# A complete map of specificity encoding enables reprogramming of a protein interaction

**DOI:** 10.1101/2024.04.25.591103

**Authors:** Taraneh Zarin, Cristina Hidalgo-Carcedo, Ben Lehner

**Affiliations:** Centre for Genomic Regulation (CRG), Barcelona Institute for Science and Technology (BIST), Barcelona, Spain; Wellcome Sanger Institute, Cambridge, UK; Universitat Pompeu Fabra (UPF), Barcelona, Spain; Institució Catalana de Recerca i Estudis Avançats (ICREA), Barcelona, Spain

## Abstract

The human genome encodes the affinities and specificities of >half a million molecular interactions. Changes in specificity drive evolution, disease and therapeutic efficacy, but how specificity is encoded and reprogrammed is not well understood. Here we present a complete map of the encoding of specificity in a protein sequence and how mutations quantitatively interact to reprogram binding. By measuring >200,000 energetic interactions we identify 17 major sites that directly and allosterically encode the specificity for six sites in a ligand. Combining mutations allows specificity for all six sites to be simultaneously re-programmed. However, binding reprogramming is not fully independent, with energetic couplings within both the protein and ligand required to accurately predict binding. This approach of measuring the specificities of thousands of proteins in a single experiment will allow many different types of molecular interaction to be understood and interpretably reprogrammed.

**Highlights:**

- Massively parallel quantification of >200,000 energetic couplings between mutations in a protein and its ligand.
- Specificity is directly and allosterically encoded in 17 sites for six positions in the ligand.
- Specificity for all six sites can be simultaneously re-programmed with accurate binding scores for >150,000 intermediately reprogrammed variants.
- Accurate binding prediction requires energetic interactions between the protein and ligand and within both the protein and ligand.

## Introduction

Specific physical interactions between proteins underlie nearly all aspects of biology from transcription and signaling to cellular trafficking, cell-cell communication and subcellular organization^1^. Eukaryotic proteomes contain hundreds of thousands of such interactions, an important class of which is globular domains binding to short linear motifs (SLiMs) embedded within intrinsically disordered regions (IDRs)^2–4^. These peptide recognition domains (PRDs) typically exist in large protein families, have affinity for many potential binding targets^5,6^, and can also be frequent targets for viral hijacking^5,7,8^. How binding affinity and specificity is encoded in these domains and the ligands they interact with is thus a major area of interest^9^ and is crucial for understanding evolution^10–14^ and human disease^15–17^, as well as protein design^18,19^ and drug discovery^20,21^.

One of the largest families of PRDs with over 270 domains in more than 150 human proteins is the PDZ (postsynaptic density 95, PSD-95; discs large, Dlg; zonula occludens −1, ZO −1) domain family^5^. These domains typically bind to IDRs at the C-termini of proteins, though they have also been found to bind internal motifs and lipids^5,6,22,23^. Historically, only two positions in the last four amino acids of PDZ ligands have been used to classify the specificity of these domains into three distinct groups^24,25^. However, more recent phage display-based studies have shown that up to six ligand positions underlie at least a dozen more specificity classes than previously considered^22^, consistent with an emerging theme that the context around motif consensus sites is important for function^17,22,26–30^.

While the last two decades have seen significant advances in high-throughput, structurally-informed, and computational studies to solve the specificity encoding problem in PRDs in general^31–34^ and the PDZ domain family in particular^12,35–37^, there are several important open questions. First, the lack of sequence conservation (∼30% sequence identity) that underlies the structurally robust fold^12^ of PDZ domains coupled with the continuous^22,35^, barcode-like^5^ specificity space has made it difficult to establish a precise link between how changes in domain sequence impact changes in specificity. Second, both PDZ domains and the ligands they bind to can be dynamic and sample different conformations: not only are there dynamic allosteric networks within PDZ domains^38–45^ that appear to be conserved^46^, but the peptides they bind to are disordered and exhibit progressively increasing motion in the bound complex away from the binding site^47^ (Fig. S2a). How these structurally heterogeneous binding modes affect the encoding of binding specificity is an open question that has yet to be addressed. Third, most studies thus far have reasonably focused experiments on residues in and around the binding site in the domain. However, no complete map of specificity-encoding exists for a PDZ domain, despite evidence that even extra-domain components can strongly affect binding via allosteric communication^27,39,43,48^. Fourth, although exceptional studies have quantified the contribution of non-additive interactions between positions in PDZ domains^13,49,50^, fewer studies have quantified the extent of additivity in positions within the ligand, where each site is assumed to be independent and represented as such in commonly used position weight matrices^51^. Indeed, the influence of non-additive interactions in the design and predictability of protein-protein interactions is an area of considerable longstanding^13,52–54^ and recent interest^55–58^ with important implications for protein engineering efforts.

Here we use deep mutational scanning^59–61^ to obtain the first complete map of how specificity is encoded in the sequence of a peptide recognition domain and how mutations quantitatively interact to reprogram binding (Fig. S1). We first comprehensively quantify the genetic and energetic landscape of peptide recognition for a model PDZ domain, including pairwise non-additive interactions between sites in the ligand. We then comprehensively mutate the 100aa domain and ligand in combination, measuring >200,000 energetic interactions that identify 17 sites in the domain that directly and allosterically encode the specificity for six sites in the ligand. Combining specificity-changing mutations in these six sites allows specificity for all six sites to be simultaneously reprogrammed. However, reprogramming of binding is not fully additive, with energetic couplings within the ligand and domain required to accurately predict changes in binding. This approach of measuring specificities of thousands of proteins by performing comprehensive mutagenesis of interacting proteins should allow many different types of molecular interaction to be understood and interpretably reprogrammed.

## Results

### Quantifying an extended genetic and energetic landscape for peptide recognition

As a model system we employed the third PDZ domain from the postsynaptic scaffolding protein PSD-95, henceforth referred to as PDZ3. PDZ3 is a class I PDZ domain, meaning it recognizes the C-terminal consensus motif X-[S/T]-X-[Φ]-COOH, where X is any aa and Φ is a hydrophobic aa^24,62,63^. Although this canonical motif involves constraint on only 2 positions in the C-terminal peptide and is sufficient for binding to PDZ3^24,64^, several studies have posited that classification of PDZ domains by class I-III is overly simplistic and that other seemingly unconstrained positions outside of and within this minimal motif affect binding affinity and specificity^5,22,47^. To comprehensively quantify the extended binding specificity of the PDZ3 domain, we measured its binding to ∼200,000 variants of a 9 aa peptide from the CRIPT protein in which the sequence of either the last four canonical (positions 0 to −3) or the preceding four aa (positions −4 to −7) were fully randomized to any of the 20 aa (Fig. 1a, Fig. S3a). In addition, we quantified binding to peptides containing all possible double aa substitutions across all eight C-terminal positions to quantify interactions between residues in the peptide (Fig. 1a).

**Figure 1.**
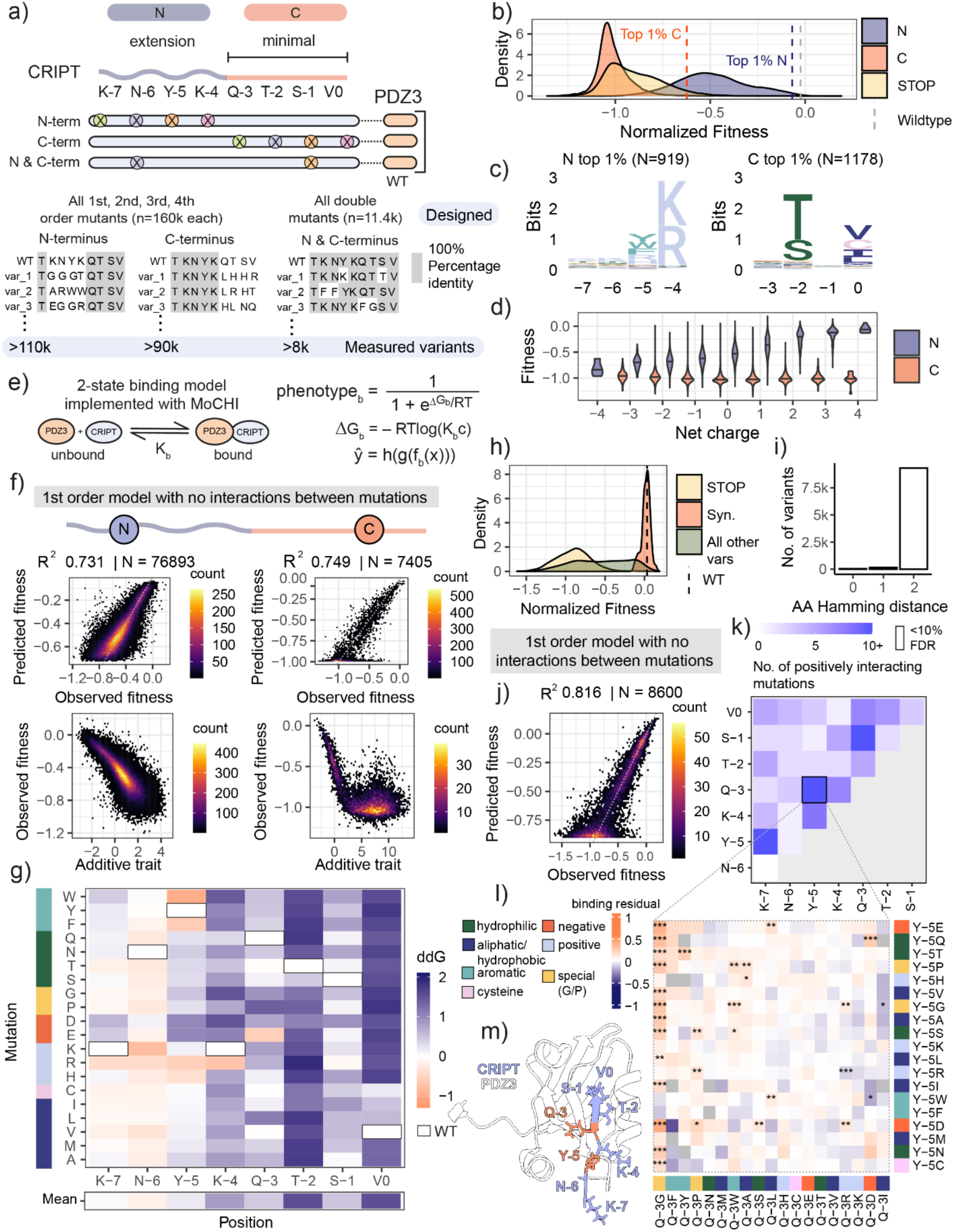
Combinatorial mutagenesis reveals energetic landscape of peptide recognition. a) Schematic showing the C- terminally bound intrinsically disordered peptide that binds PDZ3, with numbering of positions starting at the C-terminus (0) in the binding pocket to the N-terminus (-7). The C-terminal 4 residues (“minimal”) are the minimal residues required for binding, and the remaining residues (“extension”) increase binding affinity. Library design to quantify binding fitness for all possible mutants of dynamic N and structured C terminus of CRIPT as well as double mutants across the N and C terminus. In total the 3 libraries encode more than 200 000 variants in the ligand. b) Density distribution of binding fitness for N, C, and STOP variants as compared to wildtype. c) Position weight matrix (PWM) based on binding fitness scores for top 1% of variants in N vs C. d) Increasing net charge of residues in N results in incrementally increased fitness, but not in C. e) Two-state thermodynamic model implemented with MoCHI^68^ to transform binding fitness scores of mutations into energetic terms (ddG) of binding. K_b_ is the binding equilibrium constant, phentoype_b_ is the fraction bound, ΔGb is the Gibbs free energy of binding, R is the gas constant, T is the temperature in Kelvin, c is the standard reference concentration. ŷ is the output variable (predicted binding fitness), h is a linear activation function transforming the molecular phenotype to the units of the assay, g is a function for global epistasis describing the nonlinear relationship between the energetic effects of mutation and the binding phenotype, and f(x) is the additive trait map. f) Performance of model and additive trait coefficients of the N (left) and C (right) terminal positions of CRIPT. g) Heatmap of energetic binding terms for the all possible mutations (y-axis, coloured and ordered by physicochemical property) along the CRIPT peptide (x-axis). h) Density distributions of fitness and i) barplot of amino acid hamming distance for all variants for the CRIPT double mutant library. j) Performance of first order MoCHI model fit on the CRIPT cis double mutant data k) Heatmap of the number of specificity-changing mutations per pair of CRIPT positions, with one significantly enriched coupled site outlined in black (FDR<0.1) l) Heatmap of residuals for each pair of mutations in the significantly coupled CRIPT sites Q-3 and Y-5. Mutations on axes are coloured by physicochemical property. FDR of binding residuals is indicated by asterisks (*** <1%, ** 1-5%, * 5 −10%) m) Coupled residues in CRIPT (orange) on the PDZ3-CRIPT structure (5heb^62^). Note that the Y-5 aromatic sidechain was missing from the structure and was added in ChimeraX^91^ for visualization purposes.

We quantified binding using a highly-validated protein fragment complementation assay (bindingPCA)^65–67^ (Fig. S1, Fig. 1b). Binding scores for all peptide variant libraries were reproducible between replicate experiments (Pearson’s r between 0.4 and 0.8, Fig. S2b-j, Fig. S5a-d). When comparing shared variants between the C-terminal dataset and other datasets in this study, we obtain very reproducible (R^2^=0.875) fitness scores across the full dynamic range of binding for C-terminal variants (Fig. S2i-j) as we do for N-terminal variants (R^2^=0.937, Fig. S2g-h) and for the N and C double mutant library (R^2^=0.936, Fig. S5c-d).

For the C-terminal randomized library, the position-weight matrix for the top 1% of binders matched the reported consensus from previous studies using different experimental techniques ^24,62,63^ (Fig. 1c) and the distribution of binding fitness (Fig. 1b) and proportion of peptides binding to PDZ3 (Fig. S3c) sharply decreased with an increasing number of substitutions. Because of the exponential increase in the size of sequence space when combining mutations, there are actually more peptides with 4 aa changes that bind PDZ3 than peptides with one mutation (179 vs 73; 4.8-fold more, Fig. S3c). Position-weight matrices for peptides containing between one and four mutations show that the preference for T/S at the −2 position and hydrophobic residues (including cysteine) at the 0 position is maintained with increasing mutation order (Fig. S3c).

Randomizing the four-preceding aa in the N-terminus (positions −4 to −7) gave a different distribution of binding scores, with a gradual decrease in binding with an increasing number of substitutions (Fig. 1b, Fig. S3b). Although mutations in these four aa typically have smaller effects on PDZ3 binding than mutations in the C-terminal four aa (Fig. 1b), the region still makes an important contribution to binding affinity, with a preference for positively charged and aromatic residues at the −4 and −5 positions (Fig. 1c-d), and the combined effects of substitutions frequently being very detrimental (Fig. S3b).

Fitting a thermodynamic model to each dataset using MoCHI^68^ (Fig. 1e) allowed us to measure the average energetic effect of every substitution in the ligand (Fig. 1f-g). The first order model with no interactions between mutations (Fig. 1f) accounts for the non-linear relationship between the fraction bound and the Gibbs free energy of binding (dG) but otherwise assumes mutations at different sites combine additively with no pairwise or higher order energetic couplings. The resulting energy matrix predicts the experimental data very well (Fig. 1f, R^2^=0.75 and 0.73 for the C- and N-terminal randomized sequences, respectively, evaluated by 10-fold cross validation) (see Methods) and provides a complete description of the energetic effects of substitutions in all eight positions of the PDZ3 ligand (Fig. 1g). Substitutions in positions 0 and −2 are most detrimental for binding, consistent with the description of the consensus motif as X-S/T-X-Φ- COOH^23,24,62^. However, not all substitutions in positions 0 and −2 have the same energetic effects and many changes at positions −1 and −3 also cause large changes in binding energy. Moreover, mutations at position −4, which is outside of the minimal motif, have the third largest energetic effects and multiple substitutions in positions −5 and −7 cause detrimental (ddG>0) energy changes (Fig. 1g).

The energy matrix also reveals that CRIPT harbors sites where many mutations are energetically favorable (ddG<0), particularly in the N-terminal 5 positions (Fig. 1g). For example, E is energetically favored at position −3; W, F, R and H are all favored at position −5; K and R favored at -6; and R is favored at −7 (Fig. 1g). Indeed, in the first four positions but not in the C-terminal four positions, the net charge of the peptide is the strongest predictor of binding strength among hundreds of sequence features that we tested (Spearman’s rho=0.57 for N, −0.04 for C, Fig. 1d, Fig. S3d), with positively charged peptides having higher binding scores. The ability of many substitutions in the N-terminal region of the peptide to increase affinity further emphasizes the importance of this region for binding, with the longer range of electrostatic interactions^69^ compared to other non-covalent interactions consistent with a dynamic or ‘fuzzy’ contribution^70^ to affinity and consistent with previous molecular dynamics work^47^.

To evaluate the independence of mutations in the N- and C-terminal regions of the ligand we also quantified binding to a library containing >8,000 double mutants across all eight residues (Fig. 1h- i). An additive energy model was again highly predictive of binding (R^2^=0.82 by 10-fold cross validation) (Fig. 1j, Fig. S5e-f). However, quantifying the residuals between measured and predicted binding scores identified 373 interactions between mutations in the ligand (Z-test, Benjamini-Hochberg FDR<0.1, Fig. S5g), suggesting an important contribution of genetic interactions to binding in double mutants. To identify positively interacting sites in the ligand, we defined a positive intra-peptide energetic interaction when the observed binding has a positive and significant (Z-test, FDR<0.1) residual to our additive energy model (Fig. S5i) and the observed binding score is significantly different [>2 s.d.] from the binding score distribution of STOP codon variants) (Fig. S5h-i). We then identified pairs of sites in CRIPT that are enriched for these positive energetic couplings, finding that the most frequently interacting sites are at position −3 and −5 (Fig. 1k). This pair of sites contains more than 20 significant (FDR<0.1, Z-test) positive energetic couplings (Fig. 1l), most of which are linked to a change in position −3 to a G residue. Substitution to a G at −3 changes the peptides bound by PDZ3 so that any residue that is not aromatic at the −5 position is now more strongly bound than expected (Fig. 1l). This demonstrates that changes within the minimal, structured binding site can strongly affect a residue outside the binding pocket and vice versa (Fig. 1m).

Taken together, our quantification of PDZ3 binding to >200,000 ligand sequences confirms the known C-terminal binding preference, demonstrates the importance of additional residues for binding, and suggests that although specificity is well-described by an independent sites model with additive energy changes, a full description of binding specificity involves energetic interactions between sites in the ligand.

### Quantifying >200,000 energetic interactions between a PDZ domain and its ligand

Having comprehensively characterized the binding specificity of PSD95-PDZ3, we designed an experiment to understand how the binding specificity of PDZ3 is encoded in the sequence of the domain (Fig. 2a-c, Fig. S6). To identify all mutations in the globular protein domain altering the binding specificity for each individual site in the ligand as well as the corresponding specificity change, we performed a comprehensive set of double mutant cycles^71,72^ by quantifying the binding of every mutation in PDZ3 in combination with every mutation in every position in the ligand (Fig. 2a-c). Binding scores were highly correlated across three replicate experiments (Pearson’s r>0.91, Fig. S6d) as well as with shared variants in our previous libraries (Fig. S2g-j, Fig. S5c-d), allowing us to normalize binding scores across independent experiments (Methods). Consistent with our expectation^65,73^, mutations in PDZ3 detrimental for binding are strongly enriched in the binding interface with the ligand (Fig. 2d-e).

**Figure 2.**
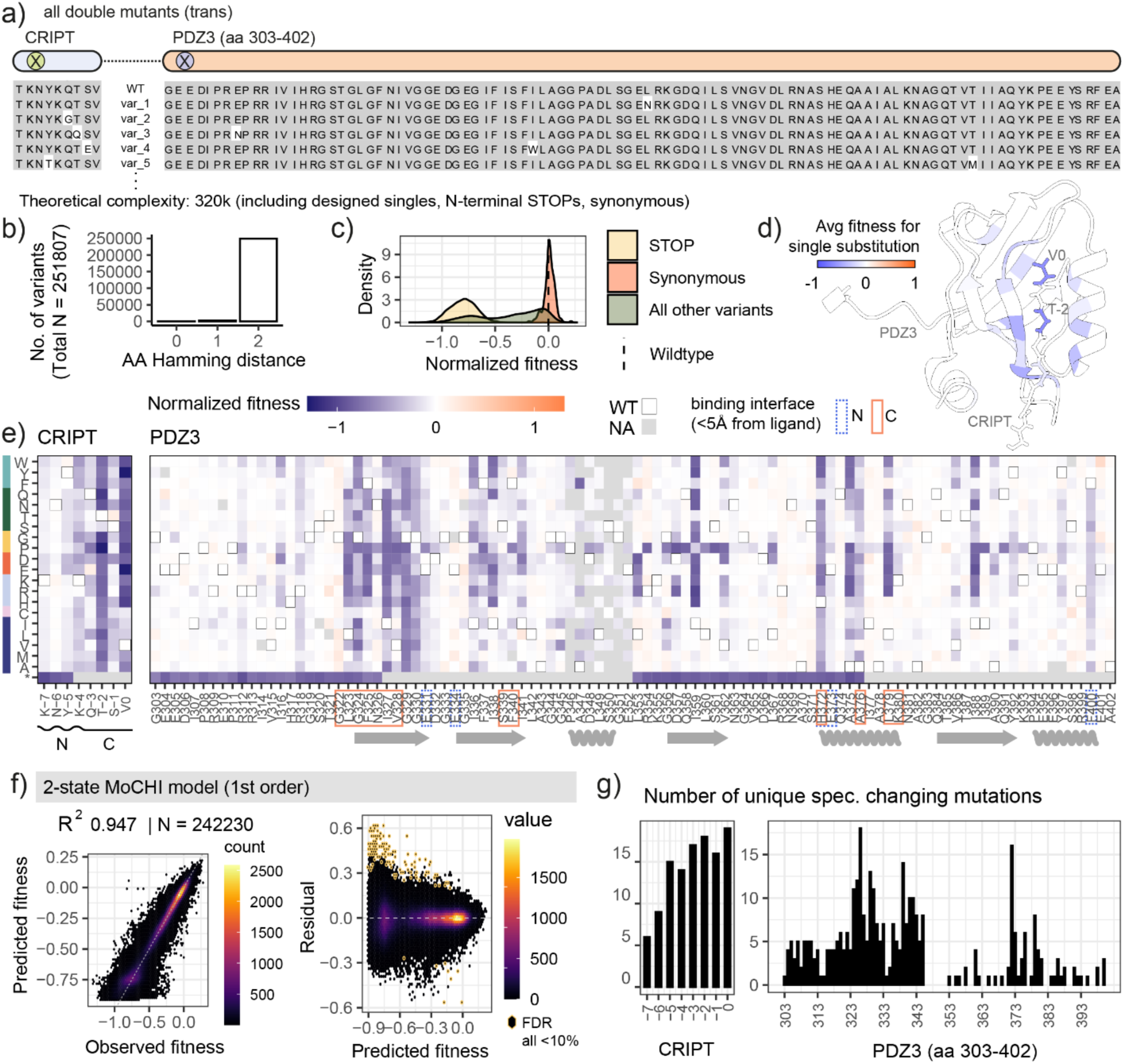
Measuring >200,000 energetic couplings between PDZ3 and CRIPT allows identification of specificity- encoding sites between the domain and peptide. a) Design of PDZ3-CRIPT trans library in which a library of all single substitutions in CRIPT is combined with a library of all single substitutions in PDZ3. Both libraries also contained designed STOP mutations and synonymous substitutions to comprise a total of 320k possible genotypes. b) AA Hamming distance and c) density distribution of binding fitness across the library for variants that pass quality thresholds. d) PDZ3-CRIPT structure (PDB ID: 5heb^62^) coloured by the average binding fitness effect for single substitutions. The canonical/constrained binding motif sites in CRIPT are labelled and have the strongest average binding fitness defects, as expected. e) heatmaps showing binding fitness of single substitutions in CRIPT (left) and PDZ3. STOP mutations are designed to only be in the N-terminus of CRIPT and two blocks of PDZ3. Most widely deleterious effects are at the binding interface (also seen in d)). f) Performance of first order 2-state MoCHI model on >240k variants (left) and residuals to the model (right) where those hexbins that pass FDR threshold of 10% (significantly different z-score from background) are outlined in yellow. g) Number of unique specificity-changing mutations (FDR<0.1) across CRIPT (left) and PDZ3 (right).

In total this dataset precisely defines the binding specificity of >1,800 globular proteins: we quantified the binding specificity of nearly every mutation in PDZ3 for nearly every mutation at every position in the ligand (Fig. S6f). Formally, changes in specificity are identified when there is an energetic coupling (or genetic interaction) between a mutation in the PDZ domain and a mutation in the ligand i.e. when binding is not well predicted by an additive energy model (Fig. 2f). We also find that the inferred free energy changes for individual mutations (ddGs, Fig. S6g) in these *trans*-double mutant libraries are highly correlated to those inferred from the N and C combinatorial ligand mutagenesis libraries, showing reproducible coefficients from modeling of independent datasets.

### Hundreds of mutations in a PDZ domain change its binding specificity

In total, we identified over 600 non-zero energetic couplings between mutations in PDZ3 and mutations in the ligand (Z-test, Benjamini-Hochberg FDR<0.1, Fig. S7a). Positive binding residuals to the additive MoCHI model (Fig. 2f) i.e. positive couplings were observed for 340 distinct mutations in 73 different positions in PDZ3 and for 114 distinct mutations in all eight residues of the ligand (Fig. 2g) (Z-test, FDR<0.1 and all binding scores significantly different from the distribution of scores for STOP codon containing variants [Fig. S7b-c]). Single aa substitutions in PDZ3 can therefore alter its specificity for all eight positions within the ligand.

### Major specificity encoding residues and the modular encoding of binding specificity

We next identified the residues in PDZ3 most important for encoding specificity for each ligand position. In total, 20 pairs of PDZ3-ligand residues are enriched for specificity-changing mutations (hypergeometric test, FDR<0.1, Fig. 3a). We refer to the PDZ3 residues within these pairs as major specificity encoding residues. There are two major specificity-encoding residues for ligand position 0, six for position −1, five for position −2, three for position −3, three for position −4, and one for position −5 (Fig. 3a, Fig. S7e). The major specificity-changing residues with the largest number of specificity-changing mutations (>20) in the domain are all physically close (<5Å distance) to the ligand position for which they change specificity (Fig. 3b).

**Figure 3.**
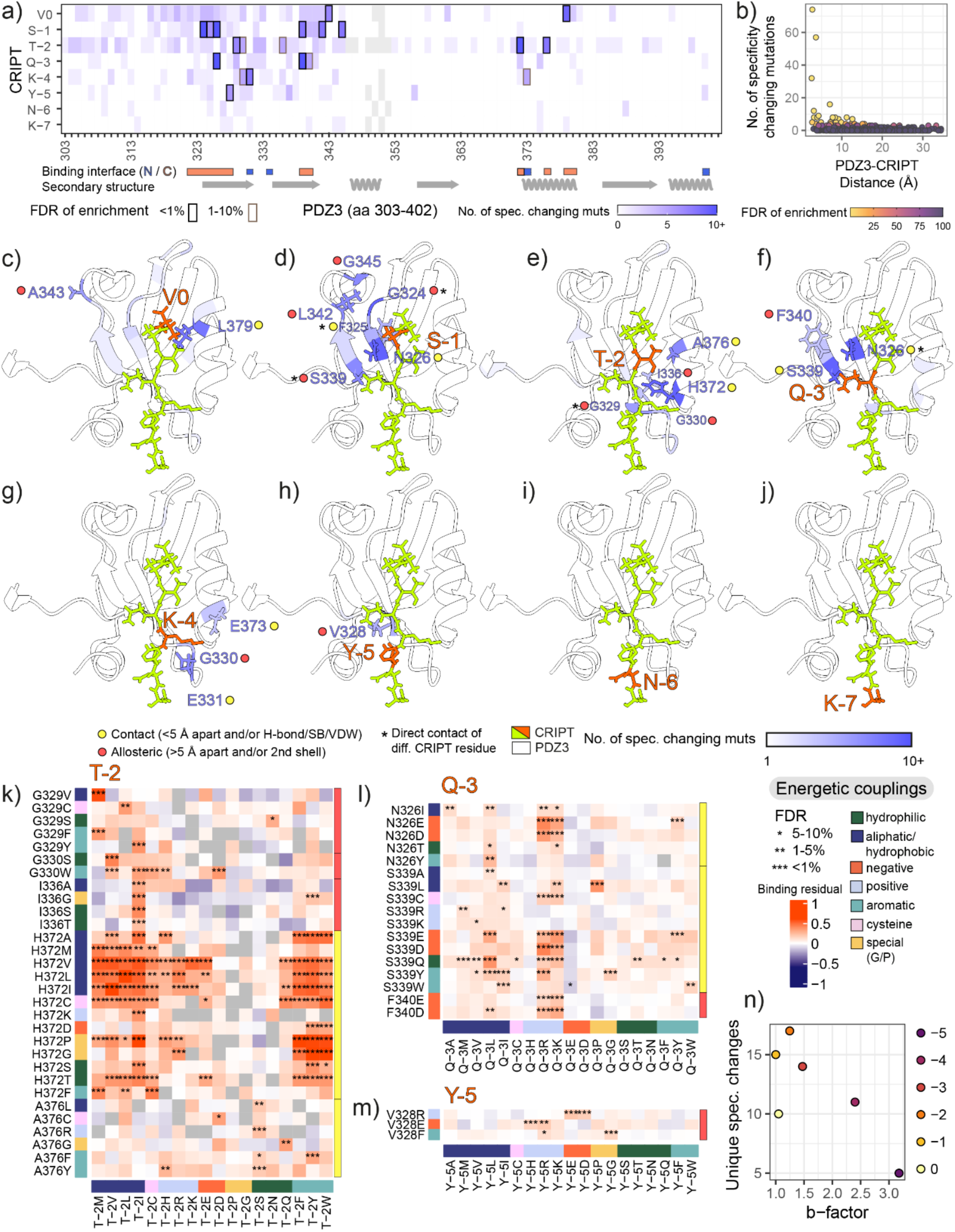
Major specificity encoding residues for each residue of a peptide. a) Heatmap showing number of specificity-changing mutations across every position-pair in PDZ3 and CRIPT. Outlined boxes mark those that pass FDR thresholds for enrichment as shown (black FDR <0.01, dark red FDR 0.01-0.1) and comprise the major specificity encoding sites. PDZ annotations for binding interface (< 5Å) and secondary structure are shown along x axis). b) Number of specificity-changing mutations across each position pair versus distance between the position pair in question. Circles are coloured by FDR of enrichment, showing as in a) that most position pairs are not enriched in specificity-changing mutations and that position-pairs that are enriched in specificity-changing mutations tend to be closer to each other in the structure. c) – j) PDZ3-CRIPT (PDB ID: 5heb^62^) structure coloured by number of specificity-changing mutations, split by the identity of the CRIPT residue (highlighted in orange [V0, S −1, T-2, Q-3, K-4, Y-5, N-6, K-7] in separate panels). Number of specificity-changing mutations is thresholded between 1 and 10 to more clearly show range for significant (FDR<0.1) sites. Only positions in PDZ3 that pass the FDR threshold of 0.1 are labelled. k) – m) Examples of specificity-changing mutations for 3 of 6 ligand positions (Full specificity map in Figure S8). Mutations are colored by physicochemical property and positions in PDZ are colored by their contact/allosteric classification for the CRIPT residue that they are coupled to. n) Number of observed unique specificity changes in CRIPT vs b-factor for each residue.

Strikingly, the major specificity encoding residues are largely distinct for each ligand position (Fig. 3a, Fig. S7d). There are 17 unique PDZ positions across the 20 pairs of positions. 14/17 PDZ positions only act as a major specificity determinant for a single ligand position, with 3 acting as major specificity determining sites for two ligand positions (Fig. S7d). The encoding of specificity in the PDZ domain is thus highly modular, with specificity for each ligand residue largely encoded by a distinct subset of residues in the domain.

The three exceptions are positions N326 and S339, which are both major specificity-encoding residues for positions −1 and −3, and G330, which defines the specificity for positions −2 and −4. N326 and S339 contact each other and −3 (N326 contacts −1 as well) (Fig. S7f-g). Similarly, G330 contacts H372 (Fig. S7g), which is a contact of position −2. G330 is also adjacent to E331, which is a contact of −4 (Fig. S7g). These positions therefore form small networks of specificity-defining residues, with clusters of PDZ3 residues connecting two separate residues in the ligand.

Eleven of these paired positions have been previously reported to be important for specificity in different PDZ domains^22,62,73^ (Fig. S7h, Table S1) but nine are novel pairs that have not been previously shown to alter specificity (Fig. S7h). Four of these nine novel pairs of sites (L379 with V0, F325 and N326 with S −1, and E373 with K-4) are contacts^24^ (Fig. S7f).

Importantly, the major specificity changing mutations that we find in the PDZ domain (293/305 mutations in major specificity encoding residues for which we obtained abundance measurements^48^) do not change *in vivo* abundance of the protein as assessed by abundance PCA (aPCA)^48,65,74^ (Fig. S7i). aPCA is a widely used protein complementation based assay that enables pooled, large-scale quantification of *in vivo* protein abundance and has most recently been applied to measure stability of over 500 human protein domains^75^, including the PSD95- PDZ3 domain that we use in this study^48,65^. When we use the abundance data in combination with binding to fit a first order 3-state thermodynamic model (Fig. S7j), the performance is nearly identical to that of fitting the binding data without abundance (R^2^=0.946 with abundance, R^2^=0.947 without abundance). Quantifying binding residuals to the model with abundance as we did with the model without abundance (Fig. 2f), we find that the energetic couplings remain nearly identical (Fig. S7k, R^2^=0.987 for >200,000 energetic couplings with and without abundance) as do the number of specificity-changing mutations (R^2^=0.994 for 784 pairs of residues, Fig. S7l). Taken together, these data strongly suggest that the major specificity changing mutations do not change the concentration of the PDZ3 domain, and that changes in PDZ3 concentration upon mutation do not explain the energetic couplings that we detect.

### Direct and local allosteric encoding of specificity

Visualizing the 20 energetically coupled pairs of residues on the structure of the PDZ3-CRIPT complex (Fig. 3c-j), shows that not only are the major specificity encoding residues distinct for each ligand position, but they are also spatially clustered. Indeed 11/17 of the major specificity encoding residues constitute the PDZ3 binding interface (Fig. 3a) and out of the 20 major energetically coupled pairs of residues, nine are direct contacts (<5Å apart and/or predicted to be contacts via hydrogen bond, salt-bridge, or van der Waals force) Fig. 3c-j, Fig. S7f), suggesting local energetic coupling as the mechanism of action.

A further 11 pairs of major specificity encoding residues are not structural contacts. These residues must therefore encode specificity either via creating new contacts or allosterically. Three of these 11 pairs involve PDZ3 residues at the binding interface that contact other residues in the ligand (Fig. 3c-j, red-filled circles with asterisks). The other eight PDZ3 residues are all contacts of PDZ3 residues that contact the ligand (Fig. S7g).

In summary, specificity is encoded both modularly and locally within the PDZ domain. Mutations in a discrete set of spatially clustered sites encode specificity for each ligand residue.

### A comprehensive map of specificity-changing mutations

For each pair of positions enriched in specificity-changing mutations, we considered how the coupled mutations in the domain and ligand relate to each other (Fig. 3k-m for three examples, full map in Fig. S8, 20x20 maps for each pair of residues in Fig. S9), as well as how energetic couplings relate to changes in binding (compared to the wild-type preference) and raw binding scores (Fig. S8). The majority of mutations that we identify in major specificity encoding residues change specificity such that the mutated ligand residue is preferred over wildtype, similar to ‘class-switching’ phenotypes^62,73^ (Fig. S8, middle panel). Others bind the mutated and wild-type ligand with similar affinity, acting similarly to ‘class-bridging’ specificity changes^62,73^.

The most diverse specificity changes (number of specificity changes out of 19 available amino acids) are observed for mutations at position −2 (17), followed closely by −1 (15) and −3 (14) (Fig. 3n). This is consistent with previous reports that found specificity “modulators” of PDZ domains at non-canonical motif positions in the ligand^76^. On the other hand, the C-terminal V0 position accommodates minimal specificity changes, a consistent result across different PDZs and experimental systems^22,77^.

Overall, we find a large diversity of mutational couplings across 6 sites in the ligand. While some positions are clearly dominated by charge-charge interactions (Y-5 with V328, Q-3 with S339, N326 and F340) (Fig. 3l-m, Fig. S7), others appear to be mediated by gain/loss of other sidechain interactions (T-2 with H372) (Fig. 3k) or more diverse changes in side-chain properties. We highlight some of the coupled sites and mutations below for the six modules defining specificity for each ligand residue.

Position −2 binding specificity is encoded by H372 (a contact of T-2) and A376 (a contact of H372 and K380, contacts of T-2 and V0, respectively) (Fig. 3k). It is also encoded by G329 (a contact of K-4 and H372), G330, and I336, all in the B2 and B3 beta-strands that contain residues directly involved in binding. Our comprehensive map finds all positions that were previously found to change specificity for T-2^22,62,73^ across multiple studies (Fig. S7h). In addition, since we have couplings for every possible amino acid, we find hundreds of new specificity-changing mutations within these sites that have not been previously identified.

The PDZ3 residue most strongly energetically coupled to position −2 is H372 (FDR<0.001, hypergeometric test, Fig. 3k). Indeed, PDZ3 position 372 and ligand position −2 are the most strongly coupled pair of positions in the dataset (table S2). This directly contacting position pair is connected by more than 70 energetic couplings. H372A was previously found to be a “class-switching” mutation^62,73^, changing the preference at the T-2 position to an aromatic residue (F). As noted in a previous study, it is not only H372A that achieves this change in specificity, but many other mutations as well^62,73^. We gain resolution into the −2 preference and find that it can also be broadened to include other aromatic residues (W/Y), as well as to hydrophobic and positively charged residues (Fig. 3k). For example, H372V/I/L/G/P/A change the position −2 specificity to aromatic, hydrophobic, and positively charged residues. H372C/T have more specific preference to aromatic or hydrophobic (but not positively charged) residues, and H372D shifts specificity to aromatic residues only. Interestingly, a fourth class of specificity change (T-2D, to an acidic residue) can be achieved with a A376C/G mutation. Based on a full hierarchically clustered map of specificity-changing residues across all sites (not just major specificity changing residues), several other mutations in PDZ can also change specificity of T-2 to T-2D (Fig. S10, blue cluster).

In terms of residues that are not direct contacts, hydrophobic and aromatic mutations in G329 change −2 specificity to hydrophobic residues (M/I/L). Similarly, mutations in the adjacent residue, G330, to V or A change specificity to I, and G330W changes specificity to I/C/V/F (Fig. 3k).

The main specificity-encoding positions for −3 are N326 (a contact of Q-3 and S −1), S339 (a contact of Q-3), and F340 (a contact of N326 and S339), meaning that the network of specificity-changing mutations around position −3 is fully connected by hydrogen bonds (Fig. 3l, Fig. S7f-g). N326 mutations to negatively charged residues (D/E) change −3 specificity to positively charged residues (R/K) and there is a similar pattern with S339, where mutations to D/E (or Q, and, more weakly, C) change preference at −3 to R/K. The same strong charge-coupling pattern can be seen for F340, where mutations to D/E create a preference for Q-3K/R.

Position −5 is the farthest coupled ligand residue from the binding pocket and has a single major specificity determinant: V328 (Fig. 3m). Once again, the couplings are charge-mediated: V328E (negatively charged) enables binding to positively-charged R/H, and V328R (positively charged) enables binding to negatively-charged D/E.

Interestingly, another residue in the extension region of the peptide that has charge-based couplings is −4 (Fig. S8). Specificity changes for position −4 occur via mutations at E331 and E373, both negatively charged direct contacts of K-4, located in the B2-B3 loop and the A2 helix, respectively, as well as G330 (a contact of E331 and shared specificity-determinant of position - 2). E331W/Y changes specificity of −4 to Y/F/W, as does G330W. In contrast, E331N changes specificity of −4 to the negatively charged residues D/E. Similarly, the polar mutation E373Q enables a −4E specificity change. The interactions between position −4 and E331 are interesting, as these residues were found to form transient salt bridges in molecular dynamics simulations^47^ as well as through our analysis of contacts from the crystal structure (Fig. S7f).

The determinants for specificity to position −1 are comprised of N326 (a direct contact of S −1), as well as S339 (a direct contact of N326), F325 (a direct contact of V0), and farther (>5A) away in the structure are G324, G345, and L342 (Fig. S8). These positions are mainly concentrated in the B2 and B3 beta-sheets and are in the binding interface (Fig. 3a). Many mutations in N326 cause the specificity-change to W and/or F/I/L in the −1 position (Fig. S8). The broadest effect (W/M/I/L/V) is seen for N326P and N326G, both amino acids with special conformations. As with Q-3, K-4 and Y-5, there are also charge-base couplings: for example, positively charged N326R/K change specificity at S −1 to negatively charged D (but interestingly, not E).

Mutations in PDZ3 that change specificity for the most C-terminal position 0 have relatively conservative effects, consistent with previous studies^77^ (Fig. S8). For example, L379 mutated to V/I changes specificity to V0F, whereas mutating L379 to polar residues G/S/T changes specificity to V0L. However, we also find that more physicochemically-diverse changes to position 0 specificity can occur through A343 mutations to M/W, both of which enable a preference for V0D. These changes increase the binding fitness of V0D from quite detrimental in the wild-type PDZ3 context to being preferred in the mutant PDZ3 (Fig. S8). Within the full map of PDZ3 energetic couplings, there is also a small cluster of mutations that enables a specificity change to V0K: E305F, E310R, G333P, and F325T (Fig. S10).

### Combinatorial quantitative reprogramming of binding specificity

Complete mutagenesis of the PDZ domain suggests a simple genetic architecture where binding specificity for six different ligand positions is encoded by six different modules of PDZ domain residues. However, in these experiments only a single mutation is made in the PDZ domain and only a single mutation in the ligand. What happens when multiple specificity-affecting mutations in the PDZ domain and/or the ligand are combined?

To address this question, we designed an experiment in which mutations in all six specificity residues of the PDZ domain and mutations in all six ligand sites are combined in all possible combinations. In the PDZ domain we included a strong specificity-changing mutation in each of the six modules (Fig. 4a) and programmed them as all single, double, triple, quadruple, quintuple and sextuple combinations (Fig. 4b-d). In the ligand we combined a total of 18 different matching specificity changes in the six positions, again as all single, double, triple, quadruple, quintuple and sextuple combinations (Fig. 4b-d). In total, there are 63 PDZ genotypes of different mutation orders paired with 2519 CRIPT genotypes of different mutation orders (Fig. S11a). Together these combine to a total of >165,000 designed genotypes including STOP and synonymous variants.

**Figure 4.**
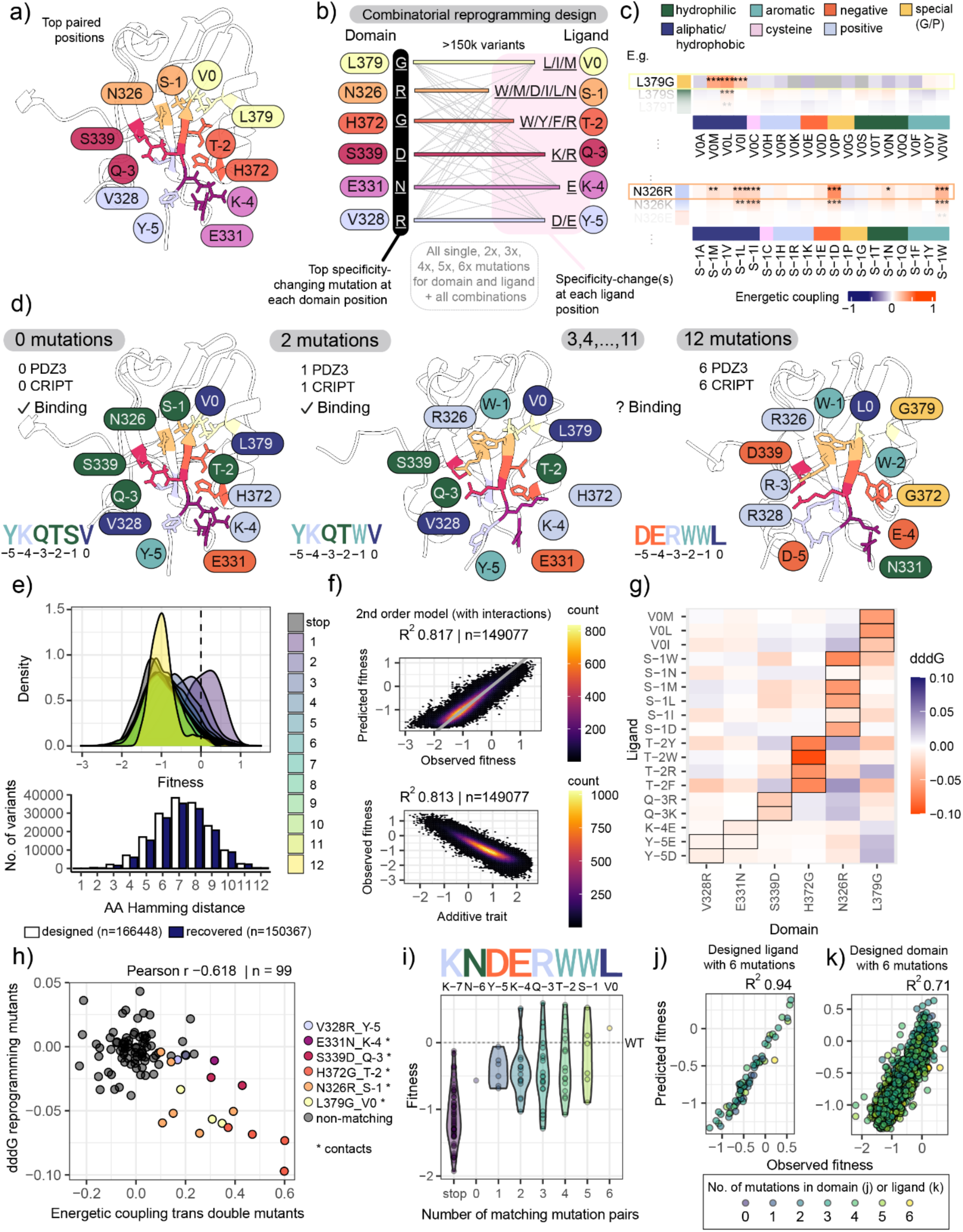
Quantitative reprogramming of binding specificity. a) Top specificity-changing positions in PDZ coloured to show how they are paired to each module/ligand position for which they change specificity in CRIPT. b) Design of reprogramming library, where the top specificity-changing mutation at each domain position is encoded along with the specificity changes it enables at each position in CRIPT. All possible single, double, triple, quadruple, quintuple, and sextuple mutations are encoded in the designed library. c) Two example specificity-changing mutations in PDZ are shown along with the mutations in the ligand that they are energetically coupled to (i.e. the specificity changes they enable shown via quantification of energetic couplings). The colour of the box around the mutation corresponds to the colour of the mutation in panel b. d) Crystal structure for wild-type PDZ3^62^ shown alongside predicted structures from ColabFold^92^ (see Methods) for PDZ3 domains with varying degrees of combined specificity-changing mutations. Physicochemical property of residue identity is coloured as per legend in panel c and corresponding ligand sequence for each structure is shown at the bottom left. Colouring of residues on structure matches panel a, denoting how the residues in the ligand and domain are coupled. e) Distribution (top) and sample size (bottom) of designed (white) and recovered (blue) library of variants with amino acid hamming distance (or stop variant) denoted by gradient of colours (purple, 1 amino acid hamming distance to yellow, 12 amino acid hamming distance). f) Performance of two-state thermodynamic model fit with MoCHI on entire reprogramming design library considering all pair-wise interactions between mutations (top), additive trait coefficients extracted from model vs. observed fitness (bottom). g) dddG (2^nd^ order energetic couplings) between PDZ specificity-changing mutations (x axis) and designed mutations in CRIPT (y axis), with corresponding specificity changes based on trans-double mutant library outlined in black. Negative dddG is a favourable interaction, blue is a detrimental interaction, units are kcal/mol. h) Linear correlation between energetic couplings inferred from trans double mutant library for all designed mutation pairs in reprogrammed library (x-axis) versus dddG energetic couplings inferred from fitting the thermodynamic model in panel f to reprogrammed library (y-axis). Dots are coloured by mutation-pair identity corresponding to panel b, and mutation pairs that are direct structural contacts are denoted by an asterisk. i) Binding fitness distribution of all variants matching one of the fully redesigned ligands (KNDERWWL) grouped by the number of mutations in PDZ3 (or STOP codon mutations in CRIPT). j) Performance of 2-state thermodynamic model in d for the designed KNDERWWL ligand with dots coloured by varying number of mutations in domain k) Performance of 2-state thermodynamic model on domain with 6 designed mutations with dots coloured by varying number of mutations in ligand.

In total we quantified the binding of 150,367 of these combinatorial mutants using bindingPCA (>90%, Fig. 4e).

Strikingly, fitting a thermodynamic model to this data shows that the binding of these combinatorial mutants is very well predicted by a simple model in which mutations have fixed energetic effects with a small contribution from pairwise energetic couplings (R^2^=0.82, Fig. 4f, Fig. S11c). Moreover, the pairwise energetic interaction terms that we quantify between the PDZ domain and ligand (dddGs, Fig. 4g) are highly correlated to the couplings we quantified from the *trans*- mutagenesis experiment (Pearson’s *r* −0.618, Fig. 4h). This indicates that the energetic couplings we quantified between pairs of mutations are robust and can be recapitulated in an independent large-scale experiment where these mutations appear in many different genetic backgrounds.

To illustrate how specificity mutations can be combined to re-programme binding to a new ligand we consider the ligand KNDERWWL. This ligand has mutations in all six specificity positions (Y- 5D, K-4E, Q-3R, T-2W, S −1W, V0L). KNDERWWL is not bound by the wild-type PDZ domain, but it is bound as predicted by the energy model by a PDZ domain with a mutation in each of the six specificity determining sites (Fig. 4i). Moreover, the energy model also accurately predicts the binding to this ligand of all 62 intermediate combinatorial genotypes i.e. the six single mutants, 15 double mutants, 20 triple mutants, 15 quadruple mutants and 6 quintuple mutants (R^2^=0.94, Fig. 4j). Similarly, the energy model accurately predicts the quantitative binding of this six-mutant PDZ domain across 2323 different ligands with between one and six different mutations (R^2^=0.71, Fig. 4k).

### Accurate prediction requires energetic couplings within the protein and within the ligand

Interestingly, accurate prediction of binding for combinatorial mutants requires not only the pairwise energetic interactions between the PDZ domain and ligand—i.e. the previously measured specificity changing mutations (Fig. 4h) — but also pairwise energetic couplings within the PDZ domain and within the ligand (Fig. 5).

**Figure 5.**
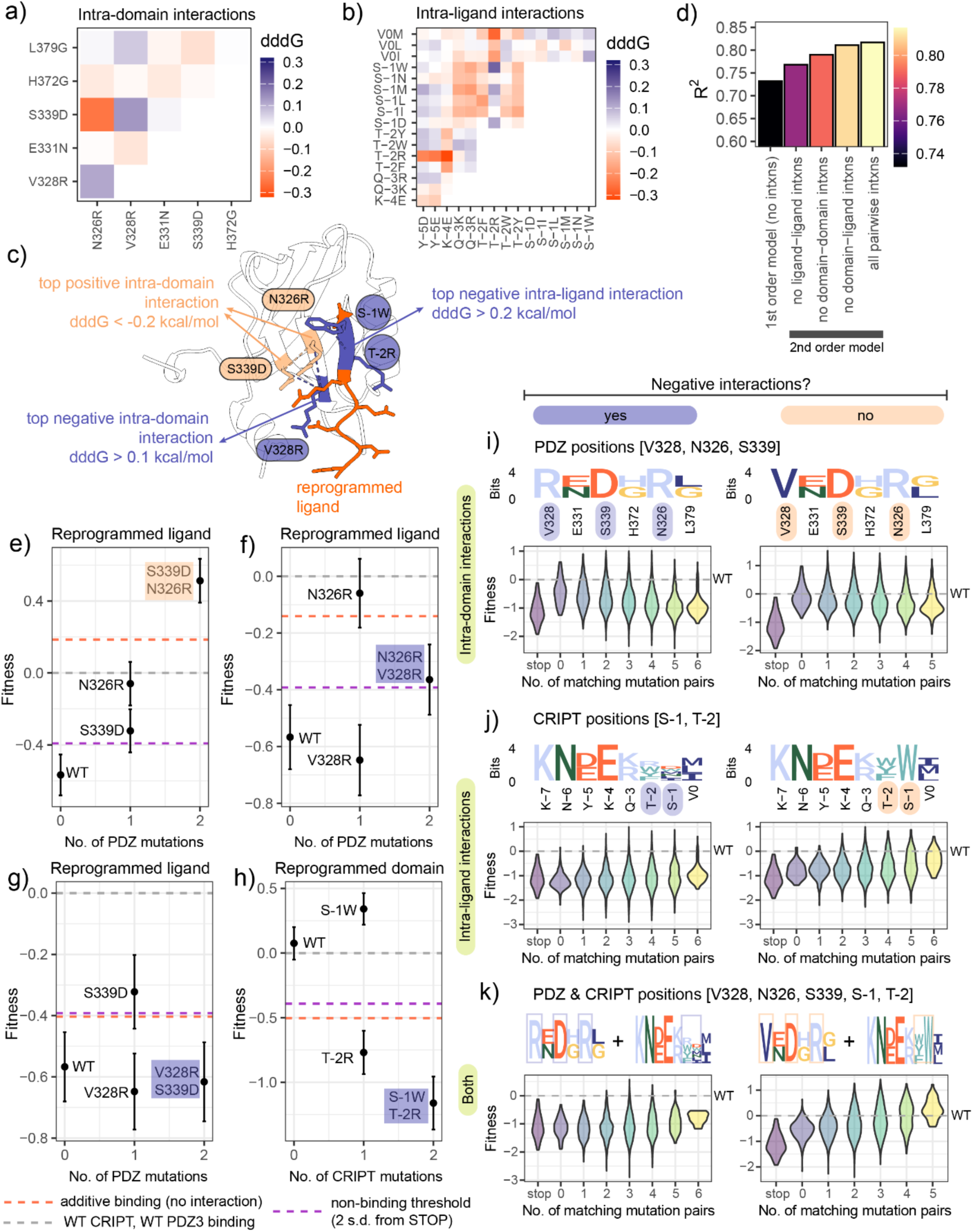
Positive and negative epistatic couplings between domain and ligand residues affect reprogramming. a) Energetic couplings inferred from 2^nd^ order 2-state thermodynamic model for intra-domain residues and b) intra-ligand residues. c) Top positive intra-domain interactions between positions 339 and 326, top negative interaction between position 328 with both 339 and 326, and top negative intra-ligand interaction between the −1 and −2 positions shown on a ColabFold^92^-derived structural model (see Methods) of the reprogrammed domain binding to the reprogrammed ligand. d) Performance of first and second order two-state thermodynamic models trained on all data (N = 149 077) considering different sets of interaction terms (i.e. coefficients) for the two-state model. e-h) measurements of binding fitness for wildtype, individual single mutants, and double mutants for variants where the two-state thermodynamic model finds epistatic interactions between ligand or domain residues. Error bars show experimental error derived from DiMSum^93^. Dashed lines show fitness values for wild-type CRIPT binding to wild-type PDZ3 (binding fitness of 0) in grey, non-binding threshold (2 s.d. from variants with a STOP codon in the ligand) in purple, and the expected fitness if there were only additive effects in the double mutant in red. i-k) Distribution of binding fitness across mutation order for all variants for which the negative interactions between domain residues, ligand residues, and combined domain and ligand residues are present (left column) or not (right column). PWM representations show which mutations are included in each position for PDZ3 and CRIPT and are coloured by physicochemical properties. Residues that have negative (purple) or positive (orange) interactions are highlighted.

With our two-state thermodynamic model, we are able to extract the pairwise interaction terms between PDZ domain residues that we reprogrammed (Fig. 5a) as well as the intra-ligand interaction terms (Fig. 5b). The strongest positive intra-domain interaction (dddG< −0.2 kcal/mol) is between N326R and S339D (Fig. 5a, c). We previously highlighted these as two of only three sites with overlapping specificity changing positions in the ligand (Fig. S7d). These coupled mutations represent two polar residues being mutated to oppositely charged residues, suggesting a charge-based mechanism for their coupling. Interestingly, the strongest negative intra-domain interaction is between the V328R mutation (another mutation to a charged residue) and both N326R and S339D (dddG >0.1 kcal/mol) (Fig. 5a, c). There is an even stronger negative interaction in the ligand (dddG >0.2 kcal/mol), between the S −1W and T-2R specificity changes. Indeed, we find that the intra-ligand interactions make the strongest contribution to the performance of the 2-state thermodynamic model that includes interactions between mutations (Fig. 5d), though the strongest performance includes all pairwise interactions compared to the first order model, resulting in a nearly 10% increase in performance.

In order to better understand how these intra-ligand and intra-domain interactions affect binding, we considered these epistatic or non-additive interactions on the binding fitness of stepwise single and double mutations on a wild-type PDZ background (binding to a reprogrammed ligand) or on a wild-type CRIPT background (binding to a reprogrammed PDZ) (Fig. 5e-h). We find that in the case of positive epistatic couplings (Fig. 5e-f), the variant with the double mutation binds better than the additive binding threshold considering no epistasis between mutations. In contrast, for those cases with negative epistatic couplings, whether in the domain (Fig. 5g) or ligand (Fig. 5h), the double mutant binds worse than expected from the additive binding threshold.

Finally, to understand how this pattern manifests in the full dataset in the space of binding scores, we quantified the distribution of binding fitness across different mutation orders including (or not) variants with negative interactions in intra-domain positions (Fig. 5i), intra-ligand positions (Fig. 5j), or both (Fig. 5k). In each case omitting negatively interacting mutations (e.g. V328R) while including positively interacting mutations (S339D and N326R) upshifted the distribution of binding fitness scores (Fig. 5i, left: mean binding fitness of bin with 6 mutations including V328R −0.95 vs. right: mean binding fitness of bin with 5 mutations not including V328R −0.34, two-sample t-test p-value <0.001). Similarly, allowing the S −1W mutation while allowing any mutation at T-2 except for T-2R also shifts up the distribution of binding fitness (Fig. 5j, left: mean binding fitness of bin with 6 mutations −0.95, right: mean binding fitness of bin with 6 mutations −0.35, two-sample t-test p-value <0.001). Interestingly, combining these effects (no negative intra-ligand or intra-domain interacting variants and including the positively epistatic variants) results in an even larger increase in the binding fitness of variants in the higher order (Fig. 5k, left: mean binding fitness of bin with 6 mutations −0.79, right: mean binding fitness of bin with 5 mutations 0.27, two-sample t-test p-value<0.001).

Overall, while there is considerable modularity and additivity in the combination of residues both in the domain and the ligand on the path to reprogramming the interaction, there are also sparse and interpretable interactions within both the protein and ligand that are important for accurately predicting binding across the >150,000 sequences.

## Discussion

To our knowledge, our data provide the first complete map of how binding specificity is encoded in the sequence of a protein domain. The map required the binding specificities of >1,800 proteins to be experimentally measured and provides a comprehensive and interpretable energy model describing how mutations throughout a protein domain reprogram its binding specificity for each site in the ligand. Combining mutations altering specificity for all six ligand sites allowed simultaneous reprogramming of specificity for all six sites and an evaluation of the energetic architecture of binding affinity changes when up to 12 mutations are combined.

In total we quantified >200,000 energetic couplings between mutations in a PDZ domain and its ligand in a pooled experiment to identify hundreds of mutations that alter the binding specificity for six different sites in the ligand. Specificity for each ligand site is primarily encoded by a spatially clustered set of residues where mutations alter specificity by a mixture of direct and local allosteric effects. Around half of the paired sites that we found to be major specificity changing residues in PDZ were novel, while the other half had previously been reported in studies using different PDZ domains.

Our data further show that modular energetic architecture allows specificity to be accurately reprogrammed by combining mutations in each of the six modules, with the binding of single, double, triple, quadruple, quintuple and sextuple mutants in the specificity-determining residues accurately predicted for both PDZ domain and ligand. The design of our combinatorial reprogrammed library allowed us to precisely measure epistatic effects in redesigned domain-ligand interactions, pointing to interesting and tractable examples of why higher order mutants can fail. Although additivity is a common and reliable assumption in the prediction of protein fitness landscapes^78^, particularly when designing stable proteins^79^, there is still much to learn about how higher order mutations affect functions such as protein binding. Our data suggests that in addition to intra-domain interactions^49^, intra-ligand interactions should be given further consideration.

Our study has a number of limitations that point to directions for future research. First, our perturbations in the trans-mutation experiment are made in the context of one specific PDZ domain binding one specific ligand. It will be important to extend this approach across the full sequence diversity of the protein family and the ligands to which they bind to gain a complete understanding of the context-dependence of specificity-encoding in PDZ domains. Second, PDZ domains are typically components of multi-domain proteins and protein complexes, with interactions between domains in the same protein and different proteins known to alter affinity and specificity^5,43,80,81^. For a complete picture of how specificity is encoded in the cell, these higher order interactions should be considered. One promising method is to perform the deep mutational scanning based complementation assays we employ here in their native cellular environments^29^.

The deep mutational scanning method that we have used here is a powerful approach for investigating how biophysical parameters are encoded in the sequences of proteins. Mutational scanning has been used to identify the residues in proteins most important for binding^12,65,67,82–84^, with titration experiments^85^ and combinatorial mutagenesis with model fitting^65,67,82^ allowing the quantification of affinity and binding energy changes for thousands of mutations in some proteins. We have applied this framework to understand the encoding of binding specificity, building on the key insight that covariation between interacting residues can reveal specificity-encoding sites^50,57,58,87^ and complementing recent mutational scanning work that quantifies how mutations in basic leucine zipper coiled-coil interactions alter their binding to other members of the family^86^.

The experimental platform that we use quantifies binding across a large dynamic range^66,67^ and is suited to quantifying the range of relatively weak interactions exhibited by PDZ domains and their targets. Weak interactions are an important part of the peptide recognition domain specificity landscape but may not be captured in the rounds of selection required for methods like phage display^5,26^, though recent developments are bridging these gaps^17,88^. Another benefit of our approach in comparison to phage display is that it does not require purification of every protein of interest and both protein and ligand sequence are genetically encoded and so easily varied alone and in combination.

The approach we have taken here can be used to understand and reprogram the binding specificity for diverse molecular interactions of varying dynamics^89^. The first step is the use of a *trans*-double mutant library to quantify how all mutations in a protein alter its specificity for each individual site in a ligand. The second step is to use combinatorial mutagenesis to simultaneously alter multiple protein and ligand sites where mutations can alter specificity. Fitting energy models to these combinatorial datasets then allows the precise quantification of the energetic couplings between and within each molecule that cause and predict the specificity changes. Provided the number of specificity-controlling residues is not too large, it should be possible to apply this approach to both protein-protein and protein-nucleic acid interactions^90^ to predictably and interpretably reprogram binding of structurally diverse proteins. With method developments, a similar strategy may also be applicable to protein-small molecule interactions to understand and interpretably reprogram protein-drug interactions.

## Acknowledgements

Funding: This work was funded by European Research Council (ERC) Advanced grant (883742), Wellcome (220540/Z/20/A), the Spanish Ministry of Science and Innovation (LCF/PR/HR21/52410004, EMBL Partnership, Severo Ochoa Centre of Excellence), the Bettencourt Schueller Foundation, the AXA Research Fund, Agencia de Gestio d’Ajuts Universitaris i de Recerca (AGAUR, 2017 SGR 1322), and the CERCA Program/Generalitat de Catalunya. T.Z. was funded by a EMBO Long-term Postdoctoral Fellowship (ALTF 525-2021) and a Marie Skłodowska-Curie Postdoctoral Fellowship (GPIDR, 101068134).

We acknowledge all members of the Lehner lab for feedback during the project, particularly Toni Beltran, Albert Escobedo, Andre Faure, Taylor Mighell, Michael Thompson, Magda Topolska, Max Stammnitz and Chenchun Weng for feedback/advice with experiments and data analysis. We thank Júlia Domingo for sharing plasmid vectors pGJJ211 and pGJJ215. We acknowledge the CRG Genomics Unit for DNA sequencing services. Molecular graphics and analyses were performed with UCSF ChimeraX, developed by the Resource for Biocomputing, Visualization, and Informatics at the University of California, San Francisco, with support from National Institutes of Health R01-GM129325 and the Office of Cyber Infrastructure and Computational Biology, National Institute of Allergy and Infectious Diseases.

## Author contributions

T.Z. and B.L. conceptualized the study. T.Z. performed all formal analysis, data curation, methodology, software, validation, and visualization. T.Z. performed all experiments except for the abundancePCA of PDZ3, which was performed by C. H-C. T.Z. and B.L. acquired funding.

B.L. provided resources and supervision. T.Z. and B.L. wrote the original draft and reviewed and edited the manuscript.

## Data availability

All raw DNA sequencing data and associated processed data files (binding fitness measurements for all datasets) have been deposited in the Gene Expression Omnibus under the accession number GSE265816. All other data required to reproduce analyses are available at https://zenodo.org/records/11048045.

## Code availability

All custom scripts required to reproduce analyses are available at https://github.com/lehner-lab/fuzzy_specificity.

## Materials and Methods

### Media

- LB: 10 g/L Bacto-tryptone, 5 g/L Yeast extract, 10 g/L NaCl. Autoclaved 20 min at 120°C.
- YPD: 20 g/L glucose, 20 g/L Peptone, 10 g/L Yeast extract. Autoclaved 20 min at 120°C.
- SORB: 1 M sorbitol, 100 mM LiOAc, 10 mM Tris pH 8.0, 1 mM EDTA.
- Filter sterilized (0.2 mm Nylon membrane, ThermoScientific).
- Plate mixture: 40% PEG3350, 100 mM LiOAc, 10 mM Tris-HCl pH 8.0, 1 mM EDTA pH 8.0. Filter sterilized.
- Recovery medium: YPD (20 g/L glucose, 20 g/L Peptone, 10 g/L Yeast extract) +0.5 M sorbitol. Filter sterilized.
- SD -URA: 6.7 g/L Yeast Nitrogen base without amino acid, 20 g/L glucose, 0.77 g/L complete supplement mixture drop-out without uracil. Filter sterilized.
- SD -URA/ADE: 6.7 g/L Yeast Nitrogen base without amino acid, 20 g/L glucose, 0.76 g/L complete supplement mixture drop-out without uracil, adenine and methionine. Filter sterilized.
- MTX competition medium: SD –URA/ADE + 200 ug/mL methotrexate (BioShop Canada Inc., Canada), 2% DMSO.
- DNA extraction buffer: 2% Triton-X, 1% SDS, 100mM NaCl, 10mM Tris-HCl pH8, 1mM EDTA pH8.

### B-factor analysis

We obtained all 214 available PDZ domain family entries bound to a ligand from the PDB. We used the Bio3D^94^ R package to read in the entries, filtering for those with 2 unique chains in the structure (n = 61), further filtered for those that had B-factors available (n = 58). We normalized the B-factors by the minimum value within that chain in order to be able to compare B-factors across crystal structures. We filtered the length of chain B to be in the first quartile of the dataset (in order to roughly filter for PDZ domains bound to a peptide ligand that was 10 aa or less) and present the 14 PDZs bound to a short ligand and their respective normalized B-factors in Fig. S2a). The names of these 14 PDZ files are listed in Table S3. The full list of 214 PDB IDs is as follows: 1gq4,1gq5,1i92,1kwa,1nf3,1nte,1obx,1oby,1obz,1r6j,1rgr,1v1t,1w9e,1w9o,1w9q,2awx,2eaq,2eg k,2exg,2f0a,2f5y,2fe5,2h2b,2h3m,2he4,2i04,2i1n,2jil,2jin,2kaw,2m0z,2m10,2ocs,2opg,2pa1,2pk t,2pnt,2q3g,2qf3,2qg1,2qku,2qt5,2r4h,2uzc,2v1w,2v90,2vrf,2vz5,2w4f,2wl7,2x7z,2zpl,3b76,3bp u,3cbx,3cby,3cbz,3ch8,3dj1,3egg,3egh,3gge,3gj9,3hpk,3hvq,3i4w,3id2,3jxt,3k82,3lgi,3lgy,3lnx, 3lny,3mh5,3mh6,3nfk,3o46,3pdv,3r0h,3r68,3r69,3tsw,3vqg,3whj,3whl,4amh,4e34,4e35,4gvc,4g vd,4h11,4jl7,4joe,4jof,4jog,4joh,4joj,4jok,4jor,4k6y,4k76,4l8n,4lmm,4mpa,4n6x,4nmo,4nmp,4nm q,4nmr,4nms,4nmt,4nmv,4nnm,4nxr,4o06,4oeo,4oep,4p0c,4p2a,4pqw,4q2n,4q2p,4q6h,4r2z,4r qy,4rr1,4uu5,4uu6,4wyt,4wyu,4xh7,4xhv,4ydp,4yyx,4z33,5d13,5e22,5elq,5ema,5eyz,5ez0,5f3x, 5fb8,5glj,5hdy,5heb,5hf4,5hff,5i7z,5ic3,5mv8,5mv9,5n7d,5n7f,5n7g,5oak,5v6t,5vwc,5vwk,5wou, 5xbf,6ar4,6bjo,6bxg,6ew9,6hks,6iqq,6ird,6jue,6lca,6lcb,6ms1,6mtu,6mtv,6mye,6nid,6q0m,6q0u, 6qjd,6qji,6qjj,6qjk,6qjl,6r9h,6rlc,6sak,6spv,6spz,6t36,6twq,6twu,6twx,6twy,6ubh,6v84,6x1n,6x1r, 6x1x,6xa6,6xa7,6xa8,6xnj,6zbq,6zbz,6zc3,6zc4,6zc6,6zc7,6zc8,7cqf,7d6f,7jo7,7lul,7ntk.

We note that the vast majority of biophysical studies of PDZ domains have been performed with minimal peptide lengths that recapitulate binding^64^. However, the dynamics of short ligands binding to PDZ domains that have thus far been used in crystallographic studies may not reflect the dynamics of the ligand when it is part of the full-length protein *in vivo*. Given the predicted disordered nature of the C-terminus and a recent Cryo-EM study of full-length CRIPT^95^, we believe it is not unreasonable to assume that the dynamics of the short ligand could be representative of the full-length ligand. However, further studies are required to understand the dynamics of C-terminal peptides bound to PDZ domains when in the context of the full-length protein.

### Library construction

We designed five libraries to probe different questions about the PDZ3-CRIPT binding interaction. All libraries had the same backbone bindingPCA vector structure of DLG4-PDZ3 (aa 303-402) N-terminally fused to DHFR3 and CRIPT (0 to -8) N-terminally fused to DHFR1,2 as in^65^. All plasmids used in the study are described in table S4. The library design for measuring PDZ3 abundance (Fig. S7i) is as described in^48^.

Combinatorial N- and C-terminal CRIPT libraries were each ordered as an NNK degenerate oligo from IDT and cloned into the bindingPCA vector containing PSD95-PDZ3 (aa 303-402) fused to DHFR3 and no bait fused to DHFR1,2 (hence “empty”). The library transformation was bottlenecked in *E. coli* such that the combinatorial space of variants was reduced from the possible 32^4^ total variants (32 possible codons at 4 positions) to 20^4^ in order to facilitate greater downstream sequencing depth for each variant.

The CRIPT “cis” double mutant library was ordered as an IDT pool of NNK oligos with 28 NNK degenerate oligos encoding all different combinations of double mutants across CRIPT positions 0 to −7. The library was transformed into the same empty vector as the CRIPT N and CRIPT C libraries described above.

The PDZ3-CRIPT “trans” double mutant library was ordered as 3 separate TWIST oligo pools. PDZ3 (aa 303-402) was split into two non-overlapping blocks for mutagenesis: Block 1 encoding the first 50 aa in the domain and Block 2 encoding the latter 50 aa in the domain. We designed 1000 variants encoding the following for each block: wildtype, all possible single mutants, half of all possible synonymous mutants, and half of all possible single STOP codon variants starting at the N-terminus of each block. Variants were designed such that each possible single mutant codon was firstly scored by the number of nucleotide substitutions away from wildtype (such that 2-3 substitutions were preferred, followed by 1, followed by 0) and then by the most optimal yeast codon (based on the *S. cerevisiae* codon usage table). This design strategy enabled us to use non-overlapping reads to sequence the long amplicons that encoded PDZ3 at one end and CRIPT at the other (schematic in Fig. S4a, designed library and results in Fig. S4b). The CRIPT single mutant library was designed in the same way but ordered as a single block due to its short length. Each library was amplified using distinct primers and transformed separately into the same backbone/bindingPCA vector (PGJJ001 as in ^65^). The PDZ3 portion of the Block 1 and Block 2 vectors were then cloned in two separate reactions into the vector with the CRIPT single mutant library to produce 2 double mutant library vectors (PDZ3 Block 1 single mutants + CRIPT single mutants, PDZ3 Block 2 single mutants + CRIPT single mutants).

The combinatorial reprogramming library was ordered as 2 separate TWIST oligo pools: one for PDZ3 and one for CRIPT. The PDZ3 pool contained all the 6 target specificity-changing mutations (N326D, V328R, E331N, S339D, H372G, L379G within aa 326-379, Fig. 4b) as single, double, triple, quadruple, quintuple, and sextuple mutations, as well as wildtype, synonymous, and STOP codon variants (at the most N-terminal position [N326] for the latter). The PDZ mutations were chosen as follows: the residual values for major specificity changing mutations were ordered first by the FDR associated with the paired position (Table S2), then sorted by residual quantile, then by abundance score quantile, and finally by residual value. The CRIPT pool contained 18 mutations at all 6 corresponding positions in CRIPT where the chosen PDZ3 mutations changed specificity (i.e. they had a positive and significant [FDR<10%] energetic coupling based on results from the trans double mutant experiment). These 18 CRIPT mutations (Fig. 4b-c) were again included as single, double, triple, quadruple, quintuple and sextuple mutations along with wildtype, synonymous, and STOP codon variants. As with the trans double mutant library, the pool of PDZ variants was combined with the pool of CRIPT mutants (all-by-all) by cloning the libraries sequentially into the bindingPCA vector (PGJJ001) to encode 63*2519 variants (Fig. S11a). We show the distribution of designed variants recovered in the assay (>90%) in Fig. 4e.

### Large-scale transformations of libraries into yeast and competition assays

We transformed each library of variants into *S. cerevisiae* in 3 replicates at a large volume scaled to the size of the library in order to ensure that all variants were present in multiple (at least 100) copies as in previous work^65,82^ to prevent bottlenecking the library. We grew the cultures to saturation in synthetic complete media with 2% glucose as a carbon source. We harvested these cultures as the “input” replicates to our selection assay, and subjected the input library to selection. The selection experiment is based on a well-described protein complementation assay^66^ wherein the yeast are grown in synthetic complete media with added methotrexate (MTX), a drug that requires dihydrofolate reductase (DHFR) for metabolization. In the presence of MTX, CRIPT (or the bait protein) must be bound to PDZ3 (the prey protein) to bring together the split fragments of DHFR and enable cell growth (Fig. S1). We harvested these cultures as the “output” from the selection assay. We extracted the DNA from each replicate using a standard phenol chloroform procedure as in^65^. To prepare the DNA for Illumina sequencing, we used two PCR steps to obtain the amplicon for each library (table S5). In PCR1, we added frameshifting oligonucleotides (table S6) and amplified the regions of interest (amplicon with constant region and mutated region) from the extracted DNA with 5 cycles. In PCR2 we added Illumina sequencing barcodes with PCR using the minimum number of cycles necessary to reach amplification plateau for each sample based on a qPCR run.

### Next Generation Sequencing and analysis of sequencing data (read counts to fitness scores)

We use sequencing as a quantitative readout for binding between PDZ3 and CRIPT (Fig. S1). We obtained reliable sequencing data for more than half a million variants across all five libraries, obtaining a fitness score (and associated error) for each variant using DimSum^93^. Parameters used to filter sequencing reads for DimSum required an input count of 10 reads in at least one replicate, and we filtered each dataset for the specific design of each library (i.e. libraries made with NNK degenerate oligos were filtered for NNK design and the custom pdz3-cript trans and combinatorial reprogramming libraries were filtered to only keep designed variants).

### Normalization of fitness scores across experiments

The CRIPT N and C combinatorial, double mutant CRIPT, and PDZ3-CRIPT trans mutant libraries contained overlapping variants that had highly correlated fitness (binding) scores when processed independently (Fig. S2g-i, Fig. 5c). In order to make all scores comparable across the study, we used a linear transformation based on these highly correlated shared variants to normalize each library (Fig. S2h-j, Fig. S5d).

### Position-weight matrices

All position-weight matrices were constructed using ggseqlogo^96^ with a custom Zappo-based^97^ color scheme to mark physicochemically related amino acids.

### Calculation of physicochemical properties

We calculated features from a curated list of amino acid property scales (n=386) (http://www.genome.jp/aaindex/) as in^98^ to quantify the correlation of these scores with the binding scores for the combinatorial N and C libraries in Fig. 1. We present those features that had a | Spearman’s r | > 0.4 for either the N or C libraries in Fig. 1d and Fig. S3d.

### Modeling phenotype to free energy with MoCHI

To translate the fitness scores, which capture the phenotypic effects of mutations, into free energy terms, we used the MoCHI package^68^ to model the fitness with a two-state thermodynamic model for protein binding. Briefly, MoCHI takes as input amino acid sequences of each variant and predicts their fitness while correcting for global non-linearities (non-specific epistasis). Using the coefficients extracted from the model, we obtain the change in free energy associated with each mutation for the phenotype in question (in our case, binding). We used default parameters for a two-state model with one phenotype (binding) for all datasets, except for the model that includes abundance in Fig. S7i. For all models, we used L1 and L2 regularization with a lambda of 10^-6^. We evaluated the model using the held-out “fold” from the 10 times that the model was run on the dataset.

The CRIPT C combinatorial library contained an overabundance (>90%) of non-binding variants, so we balanced the dataset by sampling the distribution of >100k variants without replacement so that the variants with the largest distance from the peak of non-binding variants (specified by 2 s.d. away from mean of STOP codon binding distribution) had a higher probability of being sampled (Fig. S4a-b). We used a distance function based on a power-law distribution to weight the sampling probability as follows:

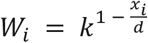

Where *W_i_* is the sampling weight of each variant binding score *x_i_*, *d* is the mean binding score for the non-binding/dead mode, and *k* is a constant (=10^10^, chosen to balance the dataset towards a reasonable percentage of binding variants [∼25%], Fig. S4b).

To test the reproducibility of the model results, we repeated the sampling procedure 10 times and performed the three implementations of MoCHI modeling (linear model with no global epistasis, two-state thermodynamic model with first order terms, two-state thermodynamic model with second order terms) (Fig. S4f). We found the modeling results to be extremely stable across 10 iterations and therefore show one representative iteration in Fig. 1f and Fig. S4c.

We note that the overabundance of non-binding variants in the C-terminal combinatorial library is also largely responsible for the comparatively lower replicate reproducibility in this dataset (Pearson’s r=0.4-0.6) compared to other datasets in this study (Pearson’s r=0.7-0.9) (Fig. S1b-c). Furthermore, while balancing the CRIPT C combinatorial library dataset is helpful for modeling first order interactions, it is detrimental for the performance of the second order model as the number of coefficients increases (Fig. S4c, f).

### Analysis of residuals to quantify energetic couplings

For both PDZ3-CRIPT trans and cis CRIPT double mutant libraries we quantified the residuals from the observed vs. mean predicted binding fitness for each variant in the respective dataset. We converted the residuals to Z-scores (using the error derived from DimSum^93^ as the denominator) and performed a Z-test to derive p-values for each variant as the Z-scores were normally distributed. We corrected the p-values for multiple testing (Benjamini-Hochberg) and report the FDR values associated with residuals where relevant. Residuals (energetic couplings) that are significant by the multiple test-corrected Z-test, >0 and pertain to variants that pass the non-binding threshold (i.e. are binding) are classified as specificity-changing. To test for enrichment of specificity-changing mutations in PDZ3-CRIPT pairs, we did a test for enrichment (hypergeometric test) of specificity-changing mutations across all pairs of sites. We again performed multiple test correction (Benjamini-Hochberg) on these to identify the major specificity encoding residues/coupled sites as those that pass an FDR threshold of 0.1. We highlight these major energetically coupled sites in Fig. 3 and list them in table S1 and S2.

### Unsupervised clustering of binding residuals

We clustered the vector of binding residuals to all possible CRIPT variants for each PDZ3 variant that had at least one specificity-changing mutation (N=340), filtered to include those PDZ3 variants with >80% data present (N=290). We used Cluster 3.0^99^ to perform unsupervised hierarchical clustering with a weighted cosine distance, average linkage and otherwise default parameters. We used JavaTreeview^100^ to visualize the full cluster plot and several manually highlighted clusters in Fig. S10.

### Protein contact determination

We used getContacts (https://getcontacts.github.io/) to predict contacting residues using get_static_contacts.py with the 5heb.pdb^62^ structure, --itypes option set to “all”, and otherwise default parameters. We also used the Bio3D^94^ package to calculate inter-residue distances between specific residues in PDZ3 and CRIPT based on the 5heb^62^ structure from PDB.

### Prediction of protein structures with multiple mutations

To predict protein structures for the PDZ3-CRIPT interaction with mutations in PDZ3 and/or CRIPT, we used ColabFold v1.5.5^92^ using template mode: pdb100 and otherwise default parameters. We show the predicted structures with 2 and 12 mutations in Fig. 4d, in Fig. 5c, and in Fig. S1 where we show the predicted structure for a reprogrammed interaction.

### Visualization of protein structures

All protein structures were based on the PDZ3-CRIPT structure with PDB ID 5heb^62^. Since the K- 7 position and the sidechain for Y-5 are missing, we added these in where relevant using ChimeraX^91^ v1.4. All represented protein structures include only those residues in PDZ3 and CRIPT that we had in our mutagenesis design, i.e. aa 303-402 in PDZ3 and 0 to −7 in CRIPT. All quantitative visualization on structures was performed with the color by attribute function in ChimeraX.

---

**Figure S1.**
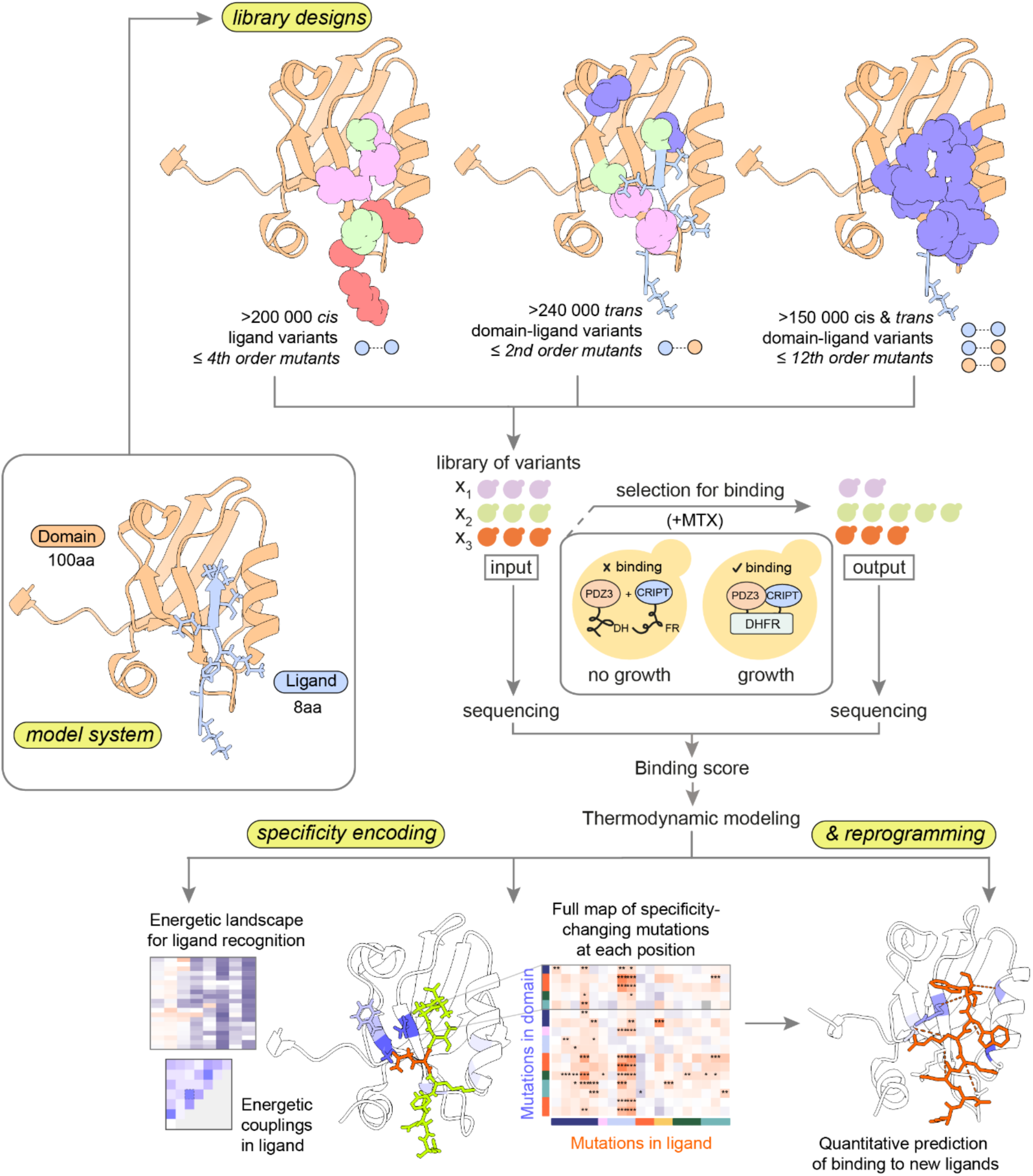
Overview of experimental design and results. Libraries of variants are transformed into yeast and selected via a protein complementation assay that quantifies protein binding^66^ (see Methods). The input library (pre-selection for binding) and the output library are sequenced and a fitness score is derived representing the enrichment or depletion of variants upon selection as compared to the wildtype sequence (see Methods). Thermodynamic modeling allows for quantification of energy terms and couplings that underlie peptide recognition and specificity encoding and allows reprogramming of the interaction.

**Figure S2.**
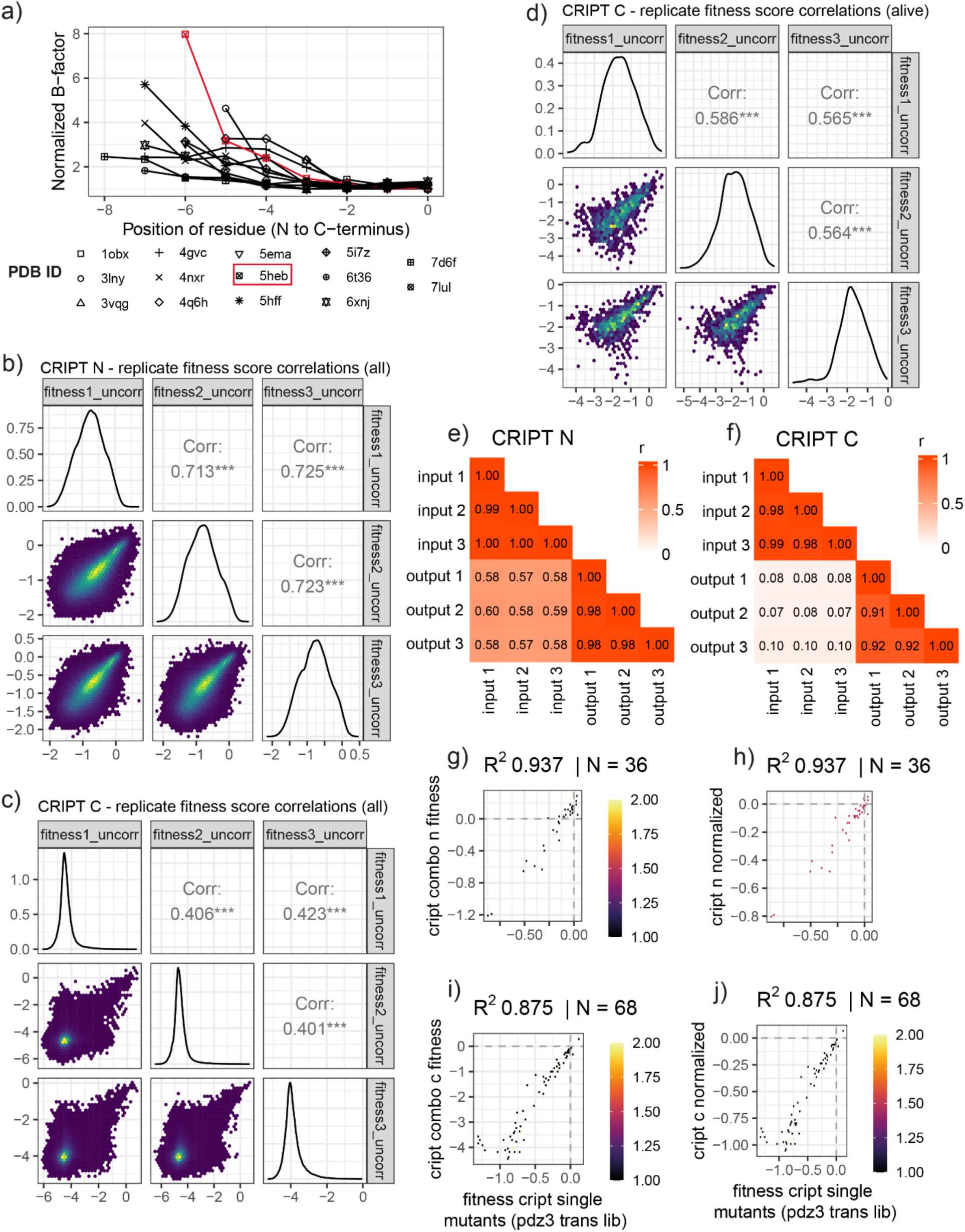
Overview of CRIPT combinatorial N and C data and quality. a) Normalized b-factor vs position of residue for PDB structures of PDZ3 crystallized with its ligand b) replicate fitness correlations (Pearson’s r) for all N-terminal CRIPT variants c) replicate fitness correlations (Pearson’s r) for all C-terminal CRIPT variants and d) top 1% of variants e) replicate count correlations (Pearson’s r) for N-terminal CRIPT and f) C-terminal CRIPT. g-i) normalization of fitness via a linear transformation using shared variants between CRIPT combinatorial libraries and the pdz3-cript trans double mutant library.

**Figure S3.**
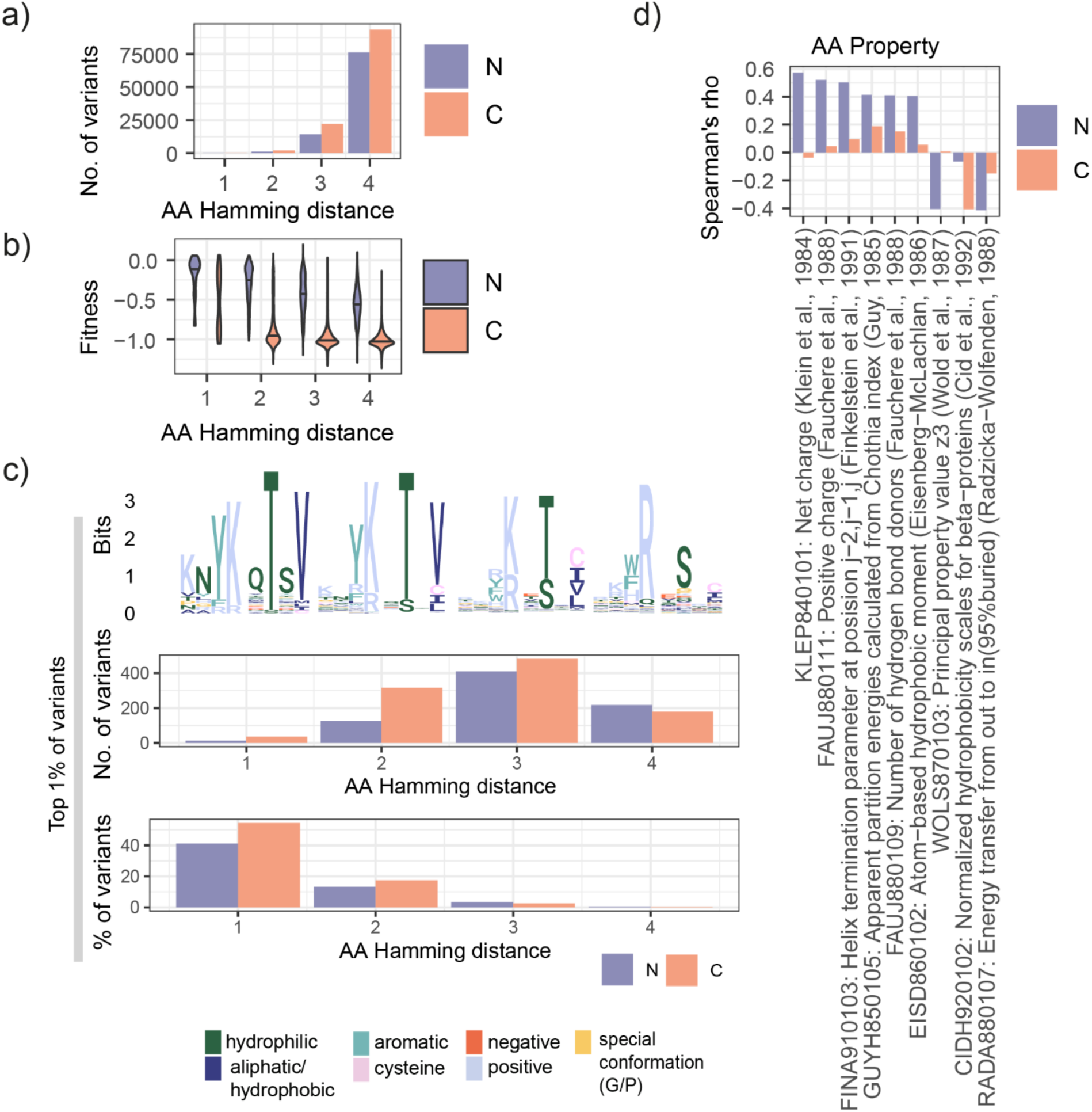
Properties of N and C combinatorial CRIPT libraries in relation to hamming distance and physicochemical properties. a) Number of variants in N vs C for each AA hamming distance away from wild-type sequence b) Hamming distance vs. fitness of N vs C libraries c) Top physicochemical features that correlate (Spearman’s rho > 0.4) with binding fitness scores in N or C dataset d) Number and percent of top 1% of variants stratified by AA Hamming distance and their associated PWMs.

**Figure S4.**
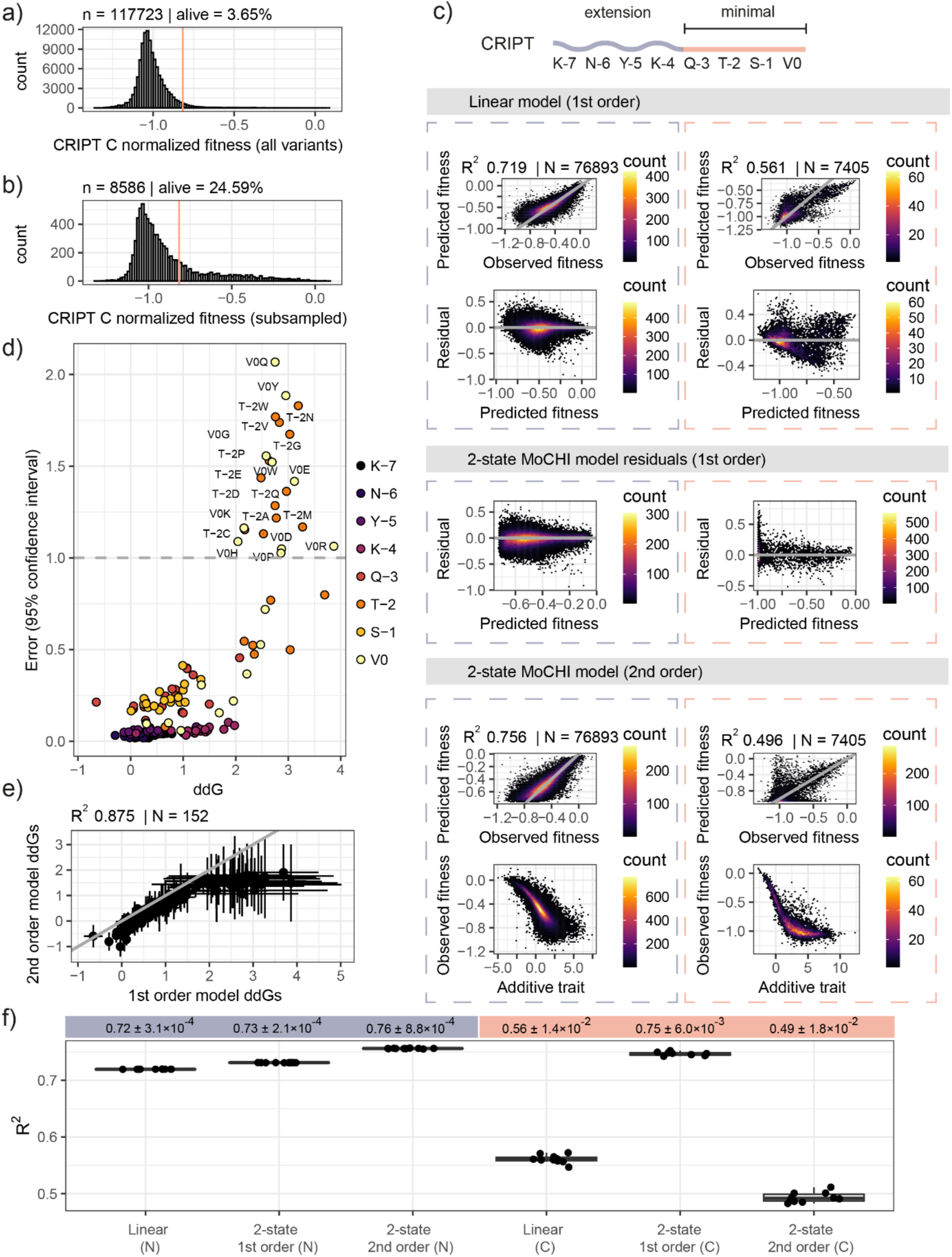
Additional MoCHI modeling results for combinatorial CRIPT N and CRIPT C datasets a) distribution of all variants in CRIPT C combinatorial library, marking percentage of variants that are “alive”/binding (i.e. 2 s.d. away from the mean of the distribution of STOP codons) b) distribution of subsampled variants that were used to balance the dataset (see Methods) c) performance of several models on CRIPT N (left) and C (right) datasets, including a linear model that does not account for global epistasis (top), residuals of a 2-state MoCHI model shown in Fig. 1 (middle), and a 2-state MoCHI model that takes into account interactions between mutations (bottom). d) The relationship between ddG values and error, with cutoff of 1 kcal/mol indicated by dashed line to represent confident ddGs – all ddG values with high error values are in the V0 and T-2 positions that also have the highest ddG values. e) correlation between ddGs from 1^st^ order vs 2^nd^ order MoCHI model for CRIPT N and C datasets. Error bars for x and y axis indicate respective 95% confidence interval in kcal/mol. f) Results from running all models on 10 iterations of subsampling the CRIPT C dataset. Mean of model performance (i.e. explained variance, R^2^ observed vs. predicted fitness) via 10-fold cross-validation for the 10 models is shown with 95% confidence intervals (2*s.d.) for each model.

**Figure S5.**
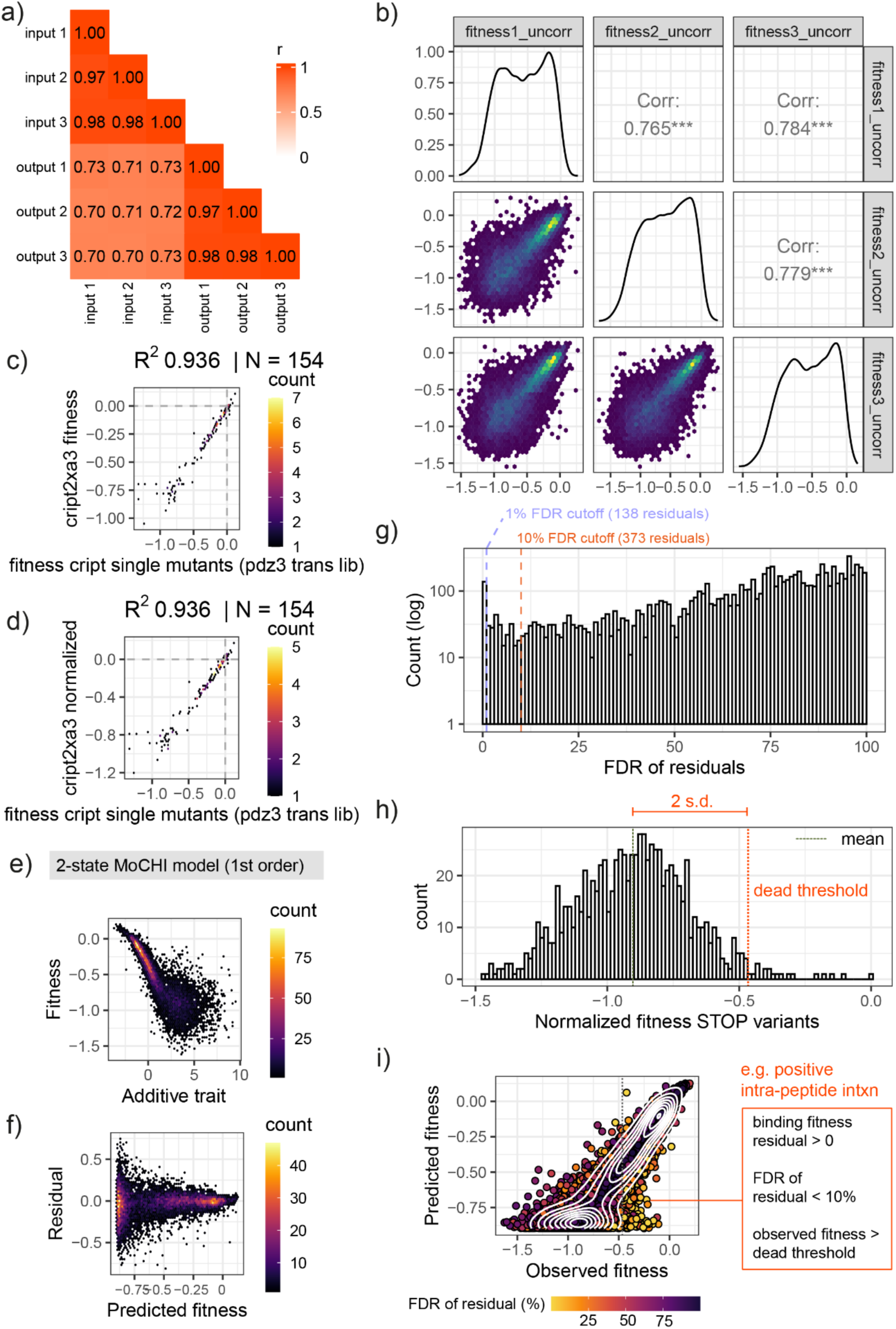
Overview of CRIPT cis double mutant library data and quality. a) replicate count correlations and b) replicate fitness correlations for all variants. c) correlation and d) normalization using linear transformation of shared variants across CRIPT cis double mutant library (y axis) and PDZ3-CRIPT double mutant trans library e) additive trait coefficients and f) residuals from fitting MoCHI two-state model g) distribution of FDR of residuals quantified from fitting MoCHI two-state model with 1% and 10% FDR cutoffs show h) distribution of variants with STOP codons and justification for non-binding/dead variants threshold (2 s.d. away from mean of STOP variants) i) Performance of two-state MoCHI model with data points coloured by the FDR of their residuals to the model fit. Dotted line shows dead threshold based on fitness distribution of STOP codon variants. Positive intra-peptide energetic couplings/interactions between mutations are demarcated.

**Figure S6.**
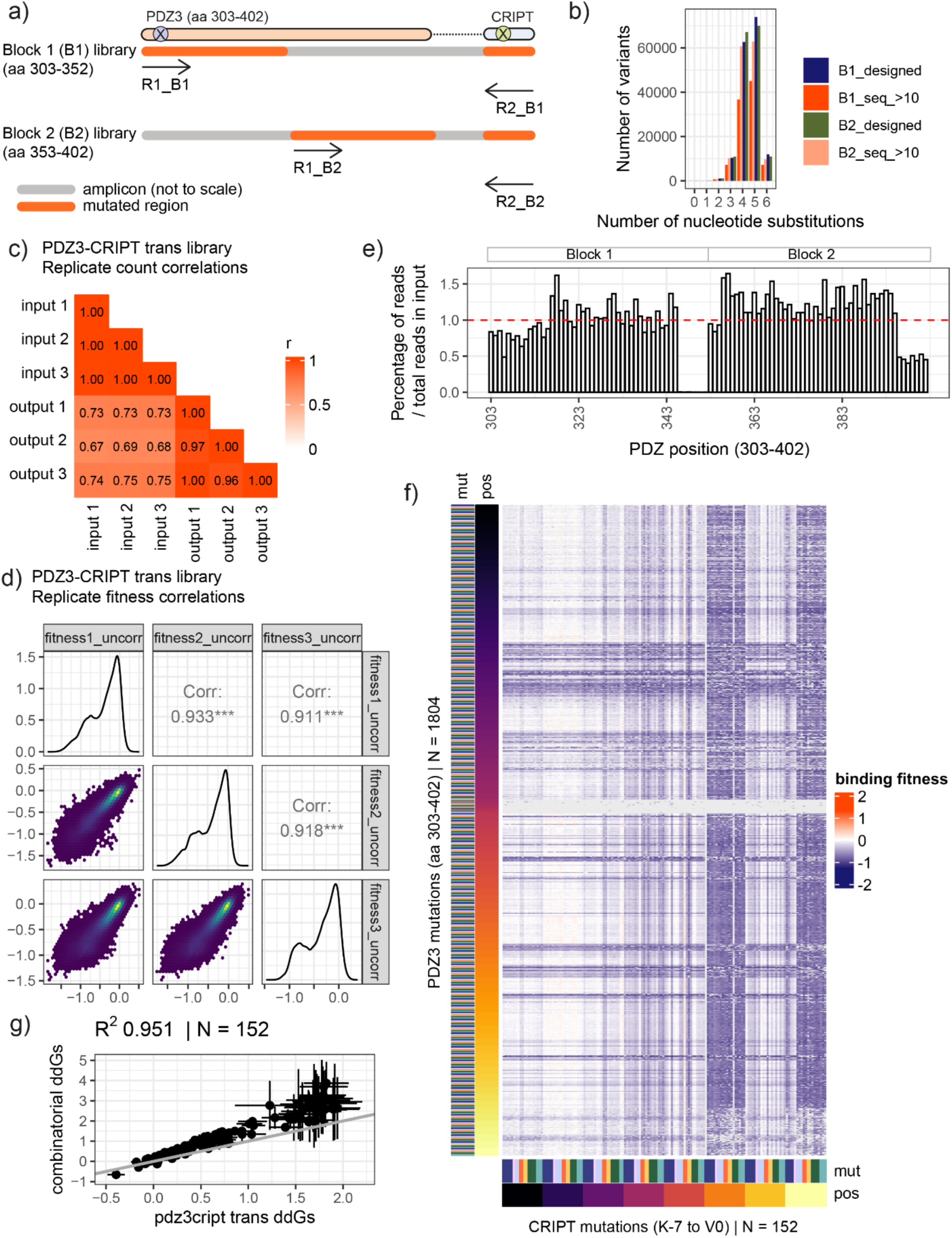
Overview of PDZ3-CRIPT trans library data and quality. a) sequencing strategy of PDZ3-CRIPT double mutant library where R1 and R2 denote forward and reverse sequencing reads, respectively. b) PDZ3-CRIPT library sequenced variants (_seq_>10 to reflect more than 10 input reads in the sample) reflect library design to maximize number of nucleotide substitutions for each encoded variant along with the most optimal yeast codon (see methods) c) replicate count correlations for PDZ3-CRIPT library d) replicate fitness correlations for PDZ3-CRIPT library e) percent of reads in input sample at each position (out of total reads in input) for each of the 100 positions assayed across the PDZ domain – even sequencing expectation is 1%, denoted by dashed red line, showing that most positions are evenly sampled except for the missing tail end of block 2. f) all-by-all heatmap of binding fitness for PDZ3-CRIPT double mutants. Mutation physicochemical properties are represented by colours as shown in Fig. 1), positions are coloured from N to C (dark to light gradient). g) correlation between ddGs from two-state MoCHI model trained on combinatorial CRIPT libraries vs. PDZ3-CRIPT libraries with corresponding 95% confidence intervals as error bars on x and y axes.

**Figure S7.**
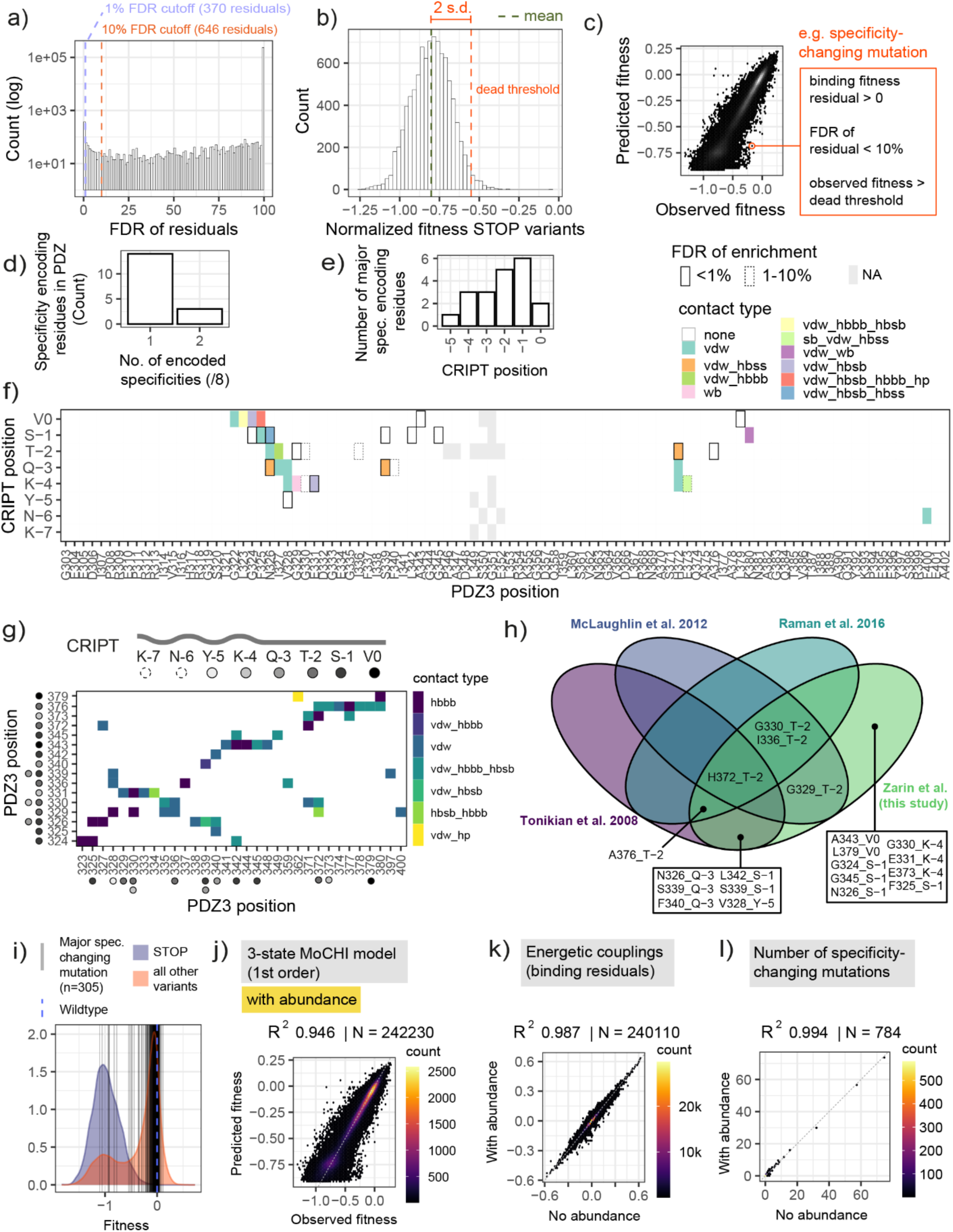
Determination of specificity-changing mutations and association with structural contacts. a) FDR of binding fitness residuals to MoCHI 2-state model for PDZ3-CRIPT mutants with different cutoffs b) distribution of STOP codon binding fitness and threshold for “dead”/non-binding threshold (2 s.d. away from STOP codon variant fitness mean). c) definition of a specificity-changing mutation d) the number of CRIPT positions for which each major specificity encoding residue in PDZ encodes specificity, showing that the vast majority encode specificity for only one CRIPT position e) Number of major specificity encoding residues (in PDZ3) for each CRIPT position (-6 and −7 have zero) f) PDZ3 position vs. CRIPT position showing position pairs enriched in specificity-changing mutations (major specificity-encoding sites) outlined by FDR value (solid vs dashed line representing <1% vs 1 −10% respectively) and the contact type of each pair as classified by getContacts (see Methods); vdw = van der waal’s, hbbb = hydrogen bond backbone-backbone, hbsb = hydrogen bond sidechain-backbone, hbss = hydrogen bond sidechain-sidechain, sb = salt bridge, wb = water bond, hp = hydrophobics g) PDZ3 positions of major specificity-encoding residues (y axis) and their contacts in PDZ3 as determined by getContacts. CRIPT residue for which each PDZ3 position encodes specificity is represented by filled circles. h) Venn diagram of all 20 major specificity encoding sites discovered in this study, and whether or not they were previously found in 3 other studies that experimentally examined change in specificity of a ligand upon mutation of a PDZ domain (see Methods for details). i) Distribution of abundance measurements for the PDZ domain (STOP= STOP codon variants, all other variants=non-STOP codon variants), with vertical dark lines indicating where the 305/323 major specificity changing mutations for which abundance measurements are available are located on the abundance fitness distribution. 293/305 of these sites are abundant (2 s.d. away from blue STOP codon variant distribution). The wildtype is indicated by a blue dotted line at 0. j) Three-state first order thermodynamic model fit to the binding measurements for trans mutations between PDZ and CRIPT as in Fig. 4.f, but also including abundance measurements k) Pearson correlation between energetic couplings (binding residuals) derived from thermodynamic model fit with and without abundance measurements included in the model l) Pearson correlation between the number of specificity-changing mutations per pair of residues with and without abundance measurements included in the thermodynamic model

**Figure S8.**
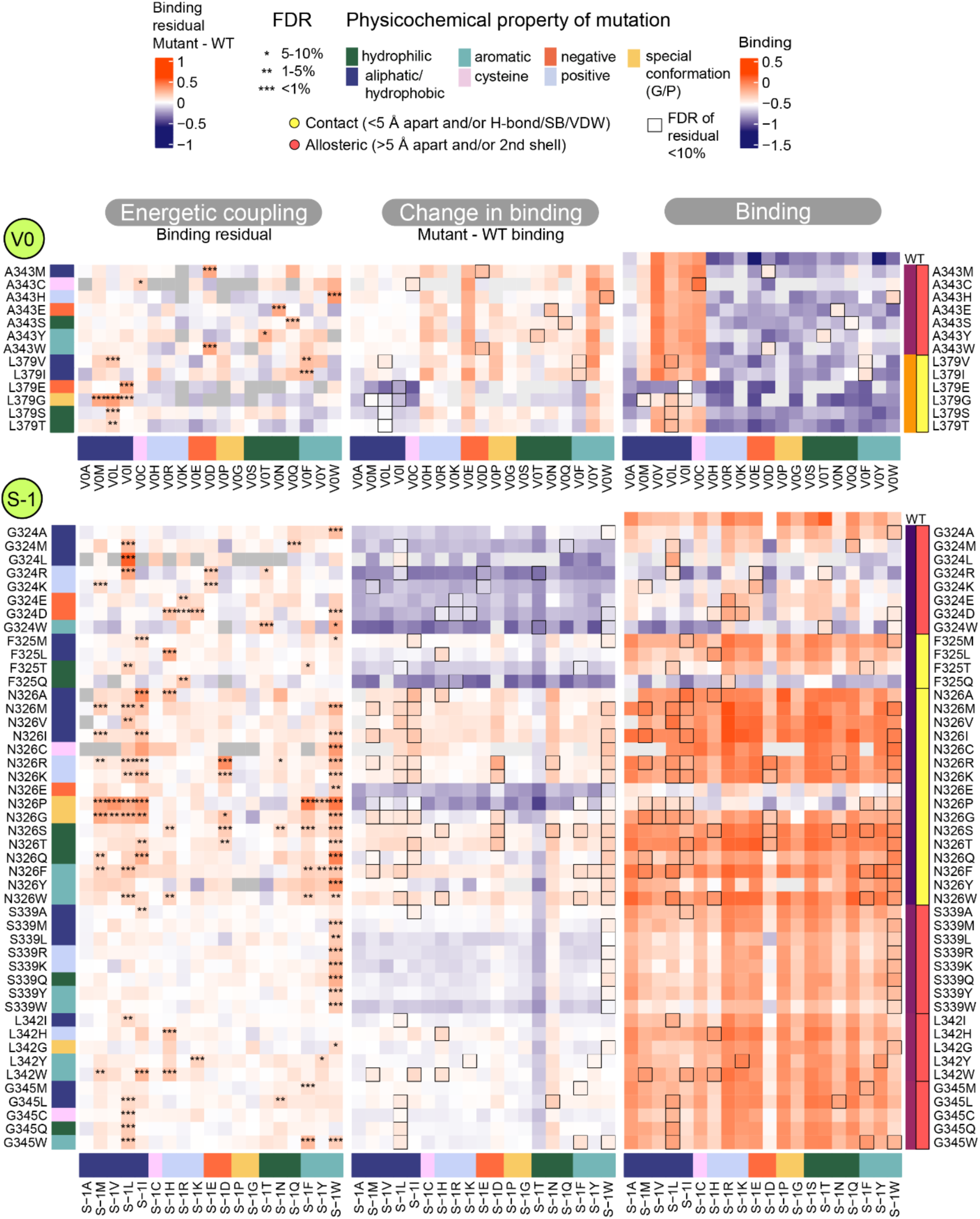
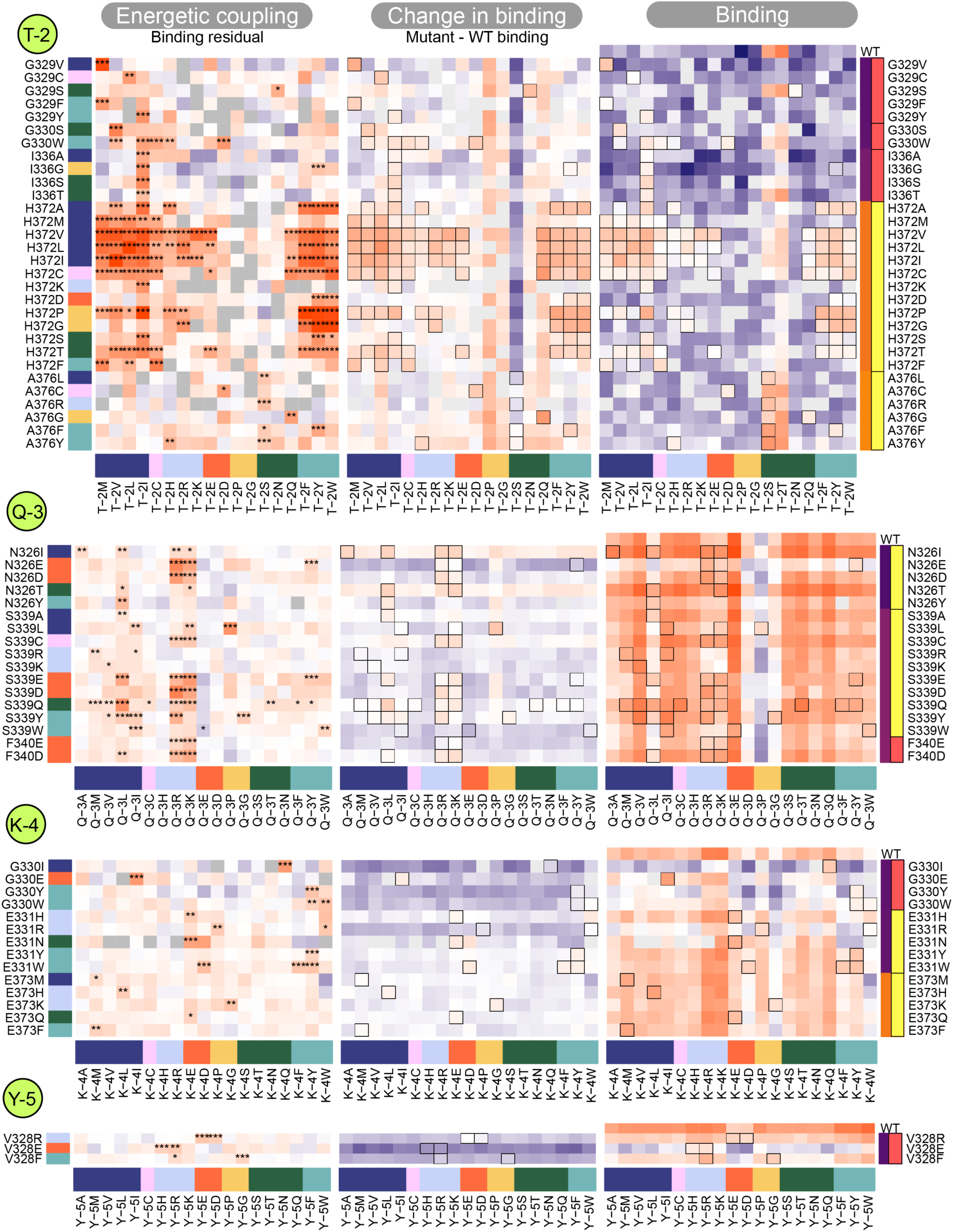
A complete map of all specificity-changing mutations across major specificity encoding sites in PDZ3-CRIPT a) Three columns of heatmaps show the energetic coupling (binding fitness residual), change in binding (binding fitness of mutant vs. wildtype), and raw binding fitness scores for all major specificity-changing PDZ3 mutations. Mutations are colored by physicochemical property and positions in PDZ are colored by their position in PDZ (from N to C-terminus) as well as their contact/allosteric classification for the CRIPT residue that they are coupled to.

**Figure S9.**
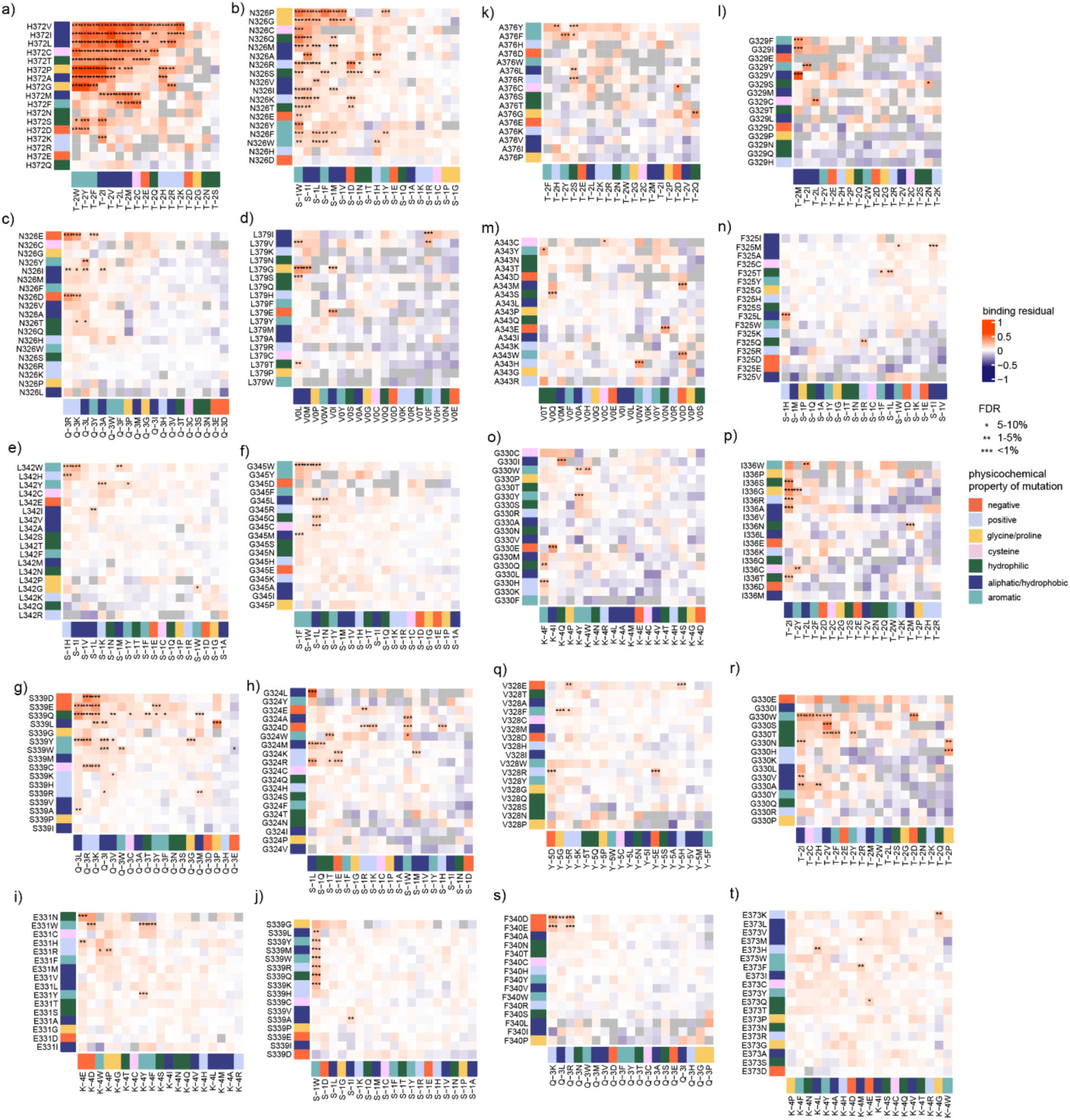
Clustered heatmaps of all 20 PDZ3-CRIPT major pairs of specificity encoding positions (i.e. significantly enriched FDR<0.1 for specificity-changing mutations).

**Figure S10.**
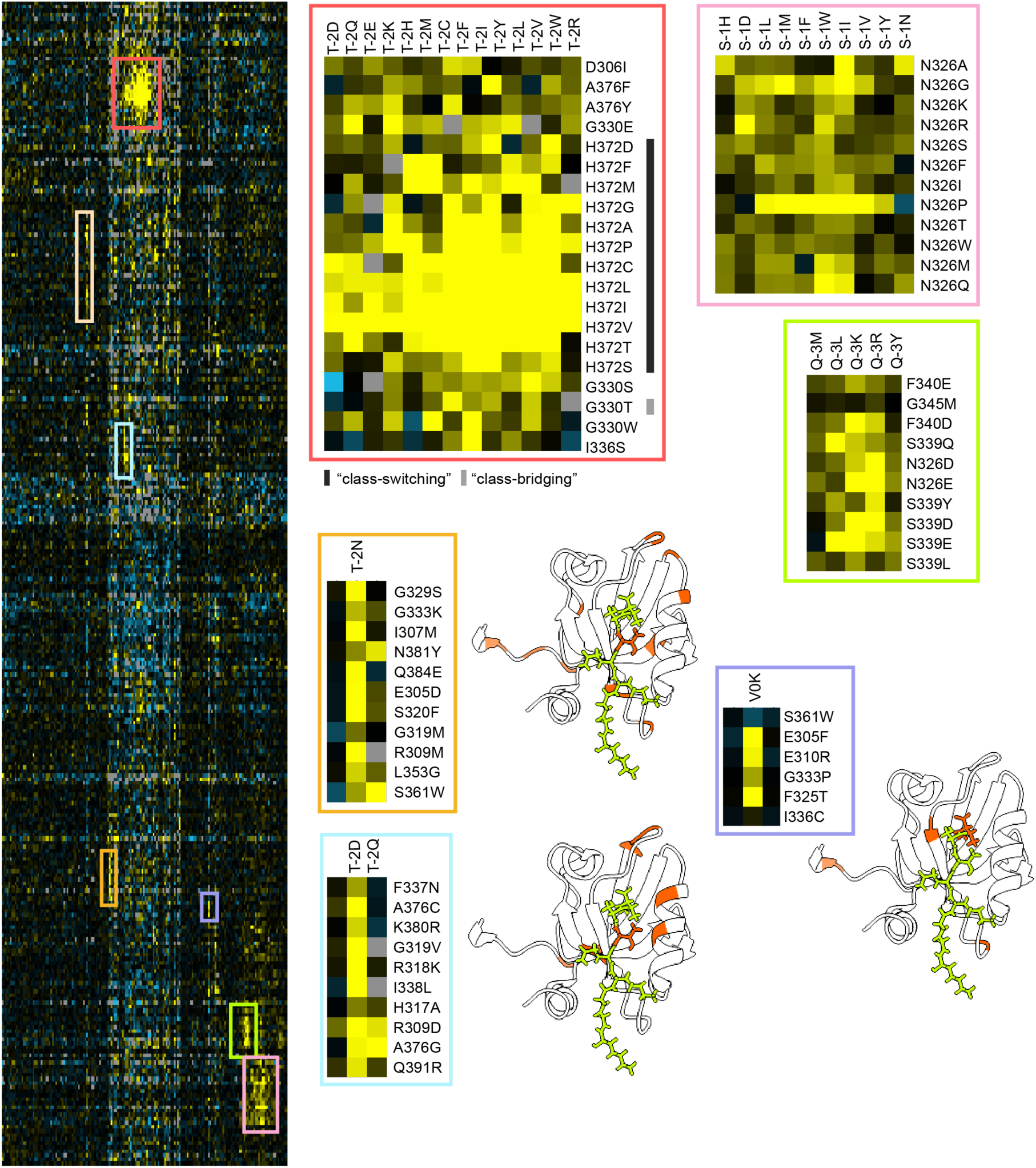
Clustered heatmap of binding residuals for all PDZ mutations (y-axis, N=290) that have at least one associated specificity-changing mutation in CRIPT (x axis, N=152). Outlined boxes represent manually-chosen clusters that point to PDZ positions enriched for specificity-changing mutations (H372, N326, S339) as shown in detail in Fig. S8 but also other sets of positions that are seemingly unrelated (orange, purple, blue clusters) with corresponding structural annotations of these sites to the right of each cluster.

**Figure S11.**
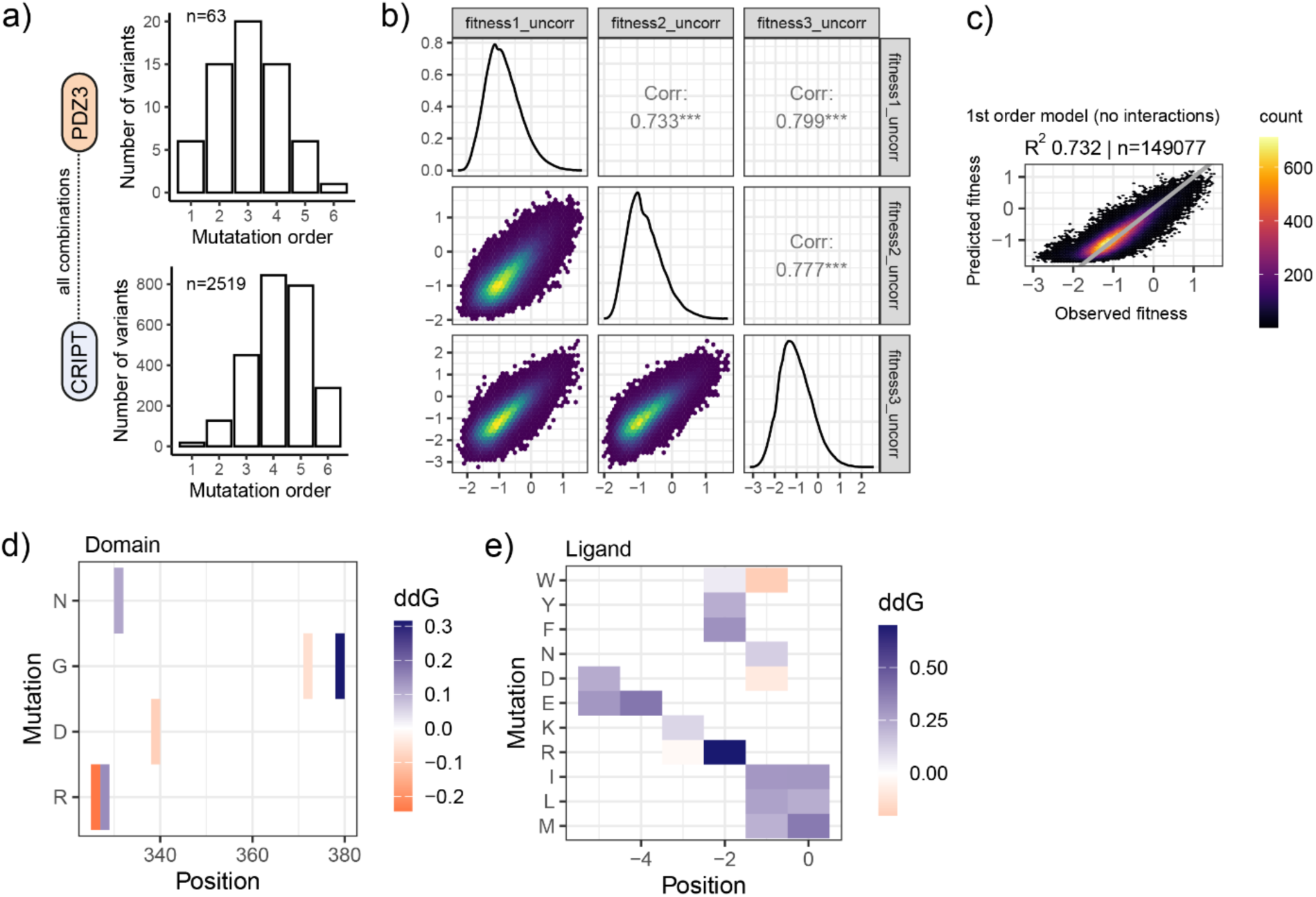
Data quality and first and second energetic terms for reprogramming library. a) library design showing number of single, double, triple, quadruple, quintuple and sextuple mutations for individual PDZ3 and CRIPT libraries that were combined to make up the reprogramming library. b) replicate correlations (Pearson’s r) for bindingPCA of reprogramming library. c) additive (1^st^ order) thermodynamic 2-state model fit using MoCHI on reprogramming library d) first order energetic terms from the 2^nd^ order two-state thermodynamic model for mutations in the domain and e) ligand

**Table S1.** Comparison of energetically coupled sites between PDZ3 and CRIPT in our study versus others that used a comparable approach. We include the positions in PDZ with standard names based on the secondary structure fold to facilitate comparison across PDZ domains.

| PDZ residue | Secondary Structure | Secondary Structure (alt) | Secondary structure class | CRIPT residue | Number of specificity changes (this study) | Major specificity change (this study) | Tonikina et al. 2008<sup>22</sup> | McLaughlin et al. 2012<sup>73</sup> | Raman et al. 2016<sup>62</sup> |
| --- | --- | --- | --- | --- | --- | --- | --- | --- | --- |
| A343 |  |  | loop | V0 | 7 | 1 |  |  |  |
| L379 | A2-8 | AB | alpha_helix | V0 | 9 | 1 |  |  |  |
| G324 | B1:B2-7 |  | loop | S-1 | 16 | 1 |  |  |  |
| F325 | B2-1 | BB | beta_sheet | S-1 | 6 | 1 |  |  |  |
| N326 | B2-2 | BB | beta_shee<br>t | S-1 | 57 | 1 |  |  |  |
| S339 | B3-4 | BC | beta_shee<br>t | S-1 | 8 | 1 | 1 |  |  |
| L342 |  |  | loop | S-1 | 8 | 1 | 1 |  |  |
| G345 | A1-2 |  | alpha_heli<br>x | S-1 | 8 | 1 |  |  |  |
| G329 | B2-5 | BB | beta_shee<br>t | T-2 | 7 | 1 |  | 1 |  |
| G330 | B2:B3-1 |  | loop | T-2 | 5 | 1 |  | 1 | 1 |
| I336 | B3-1 | BC | beta_shee<br>t | T-2 | 5 | 1 |  | 1 | 1 |
| H372 | A2-1 | AB | alpha_heli<br>x | T-2 | 74 | 1 | 1 | 1 | 1 |
| A376 | A2-5 | AB | alpha_heli<br>x | T-2 | 8 | 1 | 1 |  | 1 |
| N326 | B2-2 | BB | beta_shee<br>t | Q-3 | 12 | 1 | 1 |  |  |
| S339 | B3-4 | BC | beta_shee<br>t | Q-3 | 32 | 1 | 1 |  |  |
| F340 | B3-5 | BC | beta_shee<br>t | Q-3 | 5 | 1 | 1 |  |  |
| G330 | B2:B3-1 |  | loop | K-4 | 5 | 1 |  |  |  |
| E331 | B2:B3-2 |  | loop | K-4 | 8 | 1 |  |  |  |
| E373 | A2-2 | AB | alpha_heli<br>x | K-4 | 5 | 1 |  |  |  |
| V328 | B2-4 | BB | beta_shee<br>t | Y-5 | 6 | 1 | 1 |  |  |

**Table S2.** Major (FDR<0.1, hypergeometric test) energetically coupled pairs of sites that control specificity.

| Rank | Pair ID | CRIPT<br>N vs. C | FDR (enrichment in<br>number of spec.<br>changing mutations<br>across all pairs) | Number of<br>spec.<br>changing<br>mutations | PDZ3<br>residue | CRIPT<br>residue | Distance between<br>residues (Å) |
| --- | --- | --- | --- | --- | --- | --- | --- |
| 1 | 372_-2 | C | 2.03E-125 | 74 | H372 | T-2 | 2.715445 |
| 2 | 326_-1 | C | 1.49E-83 | 57 | N326 | S-1 | 3.593695 |
| 3 | 339_-3 | C | 1.22E-38 | 32 | S339 | Q-3 | 2.543977 |
| 4 | 324_-1 | C | 7.82E-14 | 16 | G324 | S-1 | 7.03372 |
| 5 | 326_-3 | C | 1.32E-08 | 12 | N326 | Q-3 | 3.08644 |
| 6 | 379_0 | C | 7.58E-06 | 9 | L379 | V0 | 4.245281 |
| 7 | 376_-2 | C | 3.74E-05 | 8 | A376 | T-2 | 3.924201 |
| 8 | 329_-2 | C | 0.000116 | 7 | G329 | T-2 | 9.165152 |
| 9 | 331_-4 | N | 0.000116 | 8 | E331 | K-4 | 3.126404 |
| 10 | 339_-1 | C | 0.000116 | 8 | S339 | S-1 | 8.3229 |
| 11 | 345_-1 | C | 0.000116 | 8 | G345 | S-1 | 11.18858 |
| 12 | 342_-1 | C | 0.000121 | 8 | L342 | S-1 | 7.746873 |
| 13 | 343_0 | C | 0.000607 | 7 | A343 | V0 | 12.59954 |
| 14 | 325_-1 | C | 0.007777 | 6 | F325 | S-1 | 7.572008 |
| 15 | 328_-5 | N | 0.007777 | 6 | V328 | Y-5 | 8.612266 |
| 16 | 330_-2 | C | 0.025456 | 5 | G330 | T-2 | 9.862768 |
| 17 | 336_-2 | C | 0.028273 | 5 | I336 | T-2 | 8.526768 |
| 18 | 340_-3 | C | 0.033814 | 5 | F340 | Q-3 | 4.862758 |
| 19 | 330_-4 | N | 0.057264 | 5 | G330 | K-4 | 7.177915 |
| 20 | 373_-4 | N | 0.076377 | 5 | E373 | K-4 | 2.670947 |

**Table S3.**
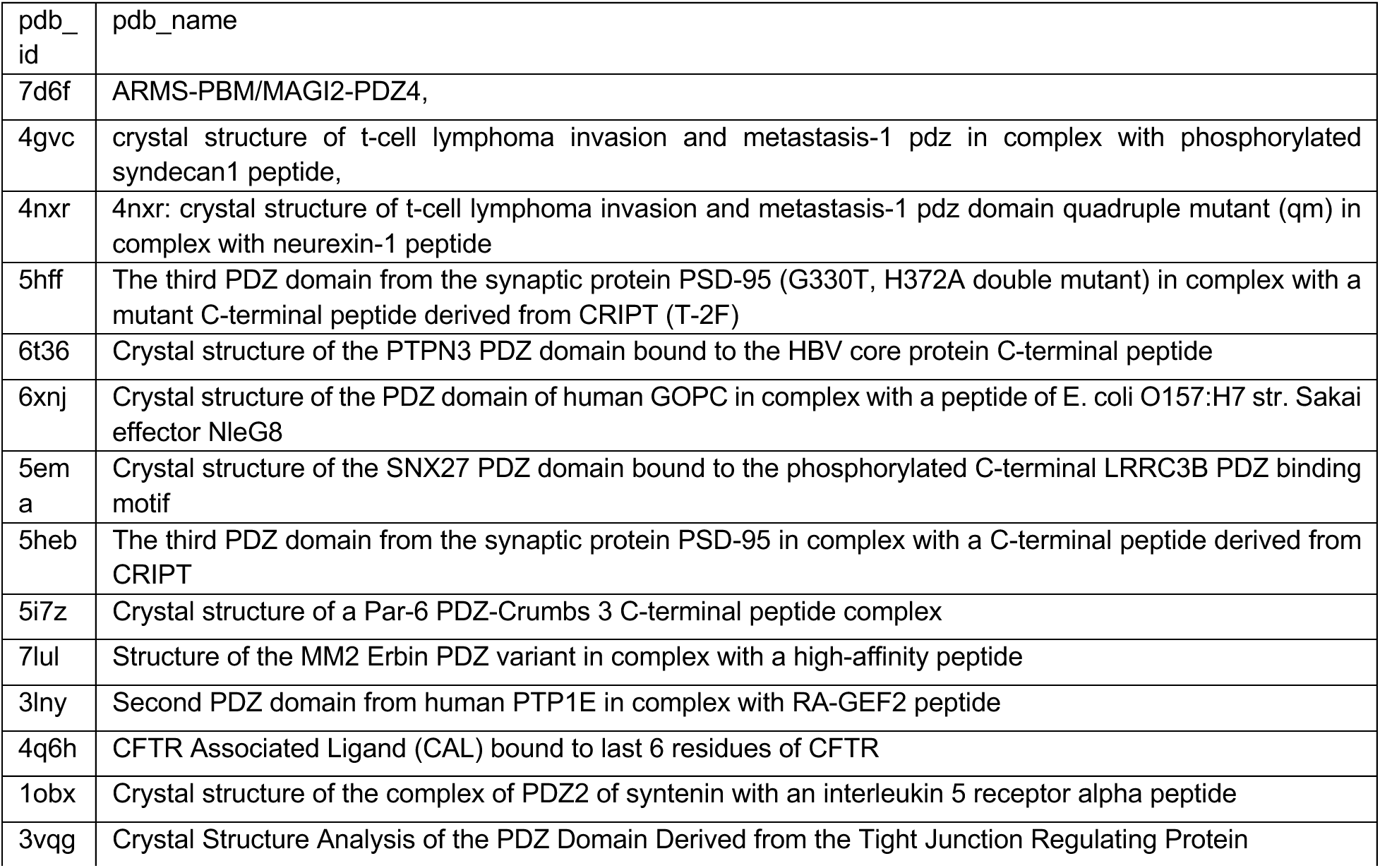
Names of PDB entries included in Fig. S2a.

**Table S4.**
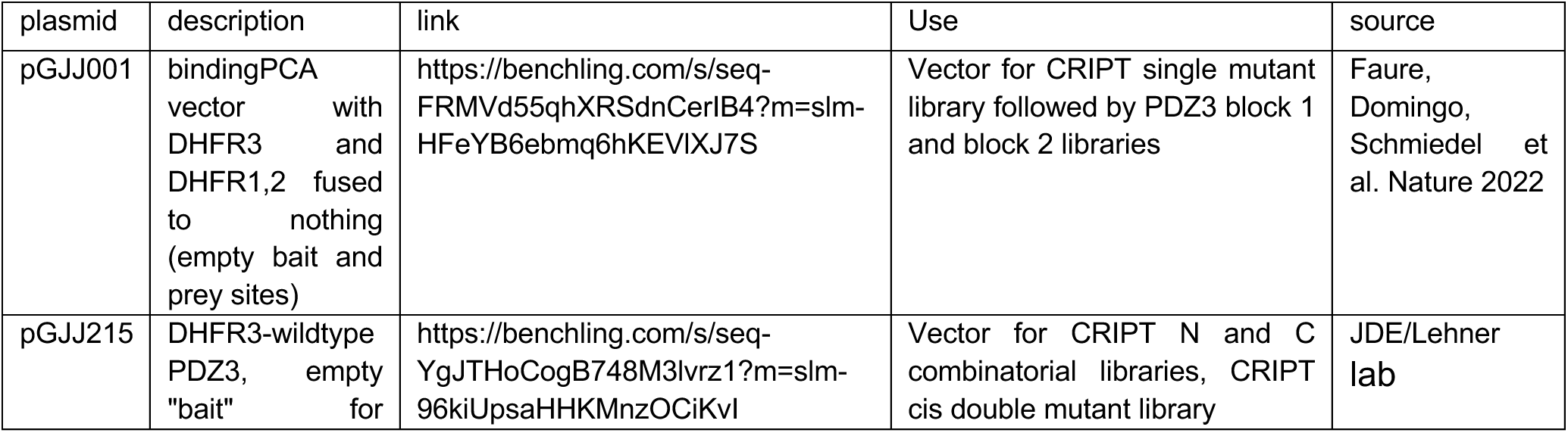
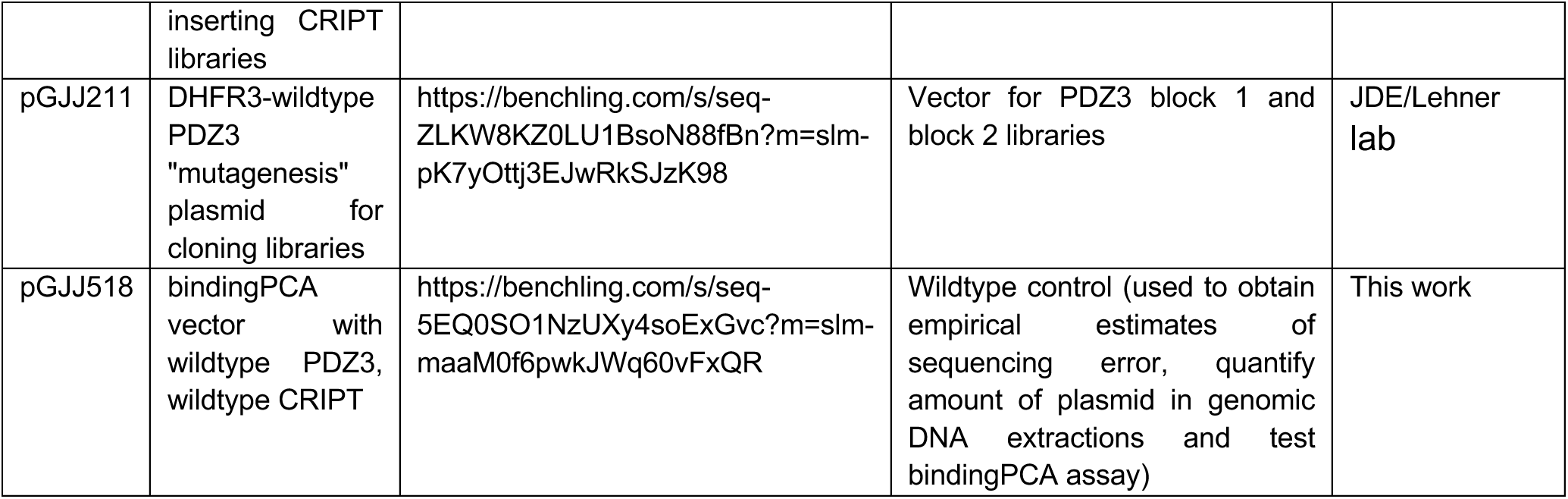
Plasmid sequences used in this study.

**Table S5.** Amplicon sequences used in this study.

| amplicon | link | notes |
| --- | --- | --- |
| CRIPT | https://benchling.com/s/seq-ZjVNfyB4q0j7ghLbVvos?m=slm-tekXiMuhgKyP7qCWdlhJ | Reference amplicon for CRIPT N, CRIPT C combinatorial libraries and CRIPT CIS double mutant library |
| PDZB1_CRIPT | https://benchling.com/s/seq-m7a9OYYNBzp1HrSv0cgQ?m=slm-j3wIHLONBYjdpZy7uZVO | Reference amplicon for PDZ3-CRIPT -- block 1 of PDZ3 |
| PDZB2_CRIPT | https://benchling.com/s/seq-5krvszGJdAW94riW4Pmi?m=slm-b0Ku0YyzgXMeTFhjQ54N | Reference amplicon for PDZ3-CRIPT TRANS double mutant library -- block 2 of PDZ3 |

**Table S6.**
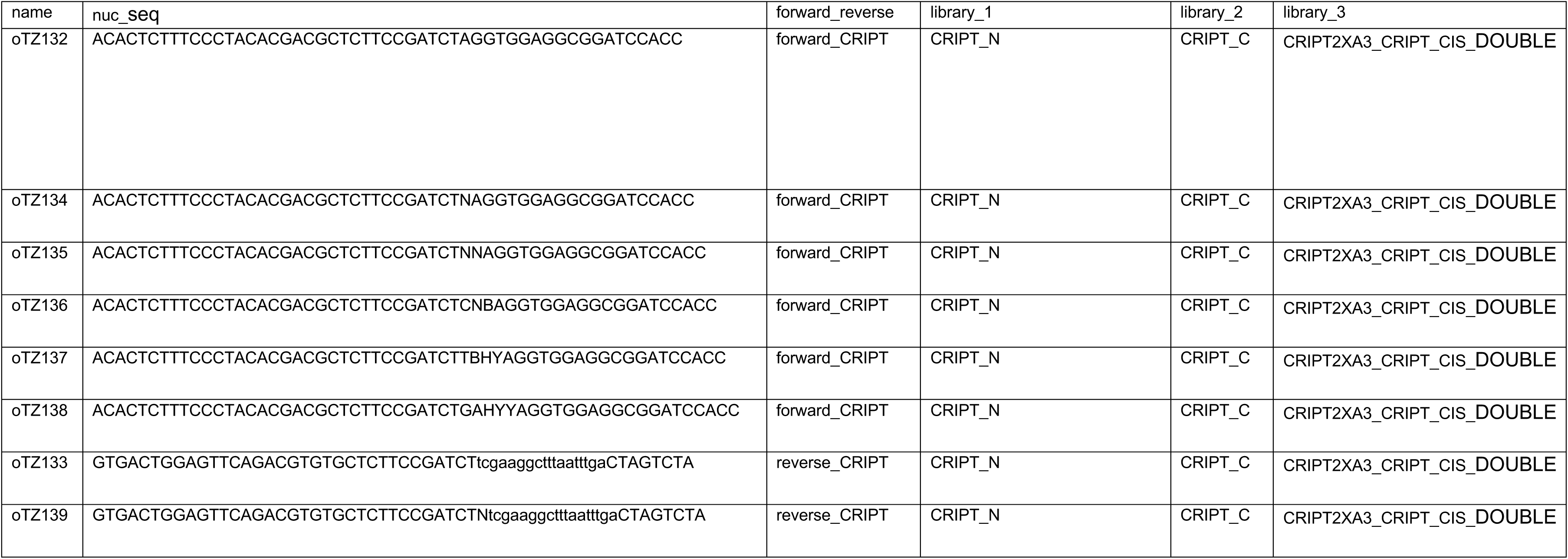
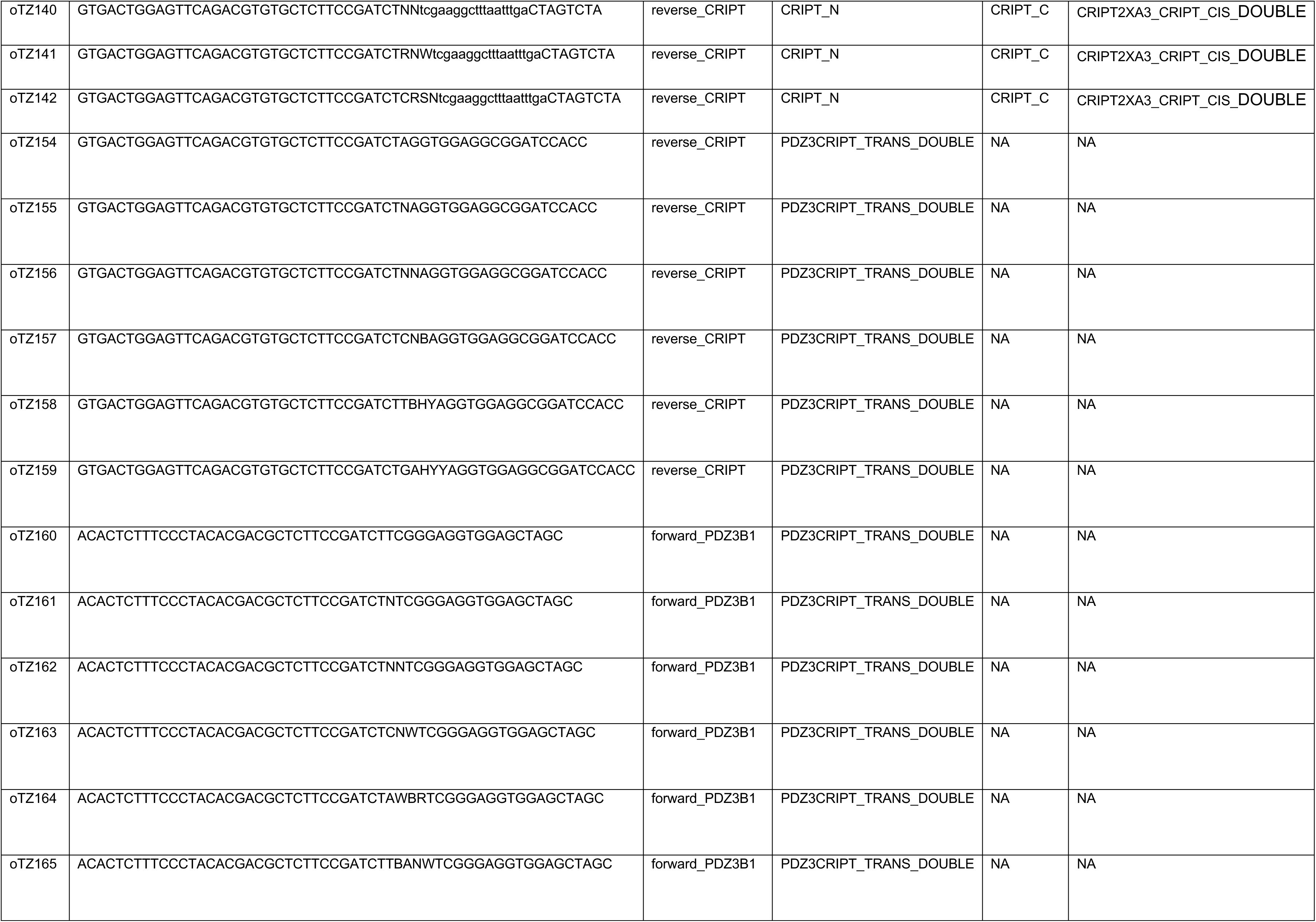
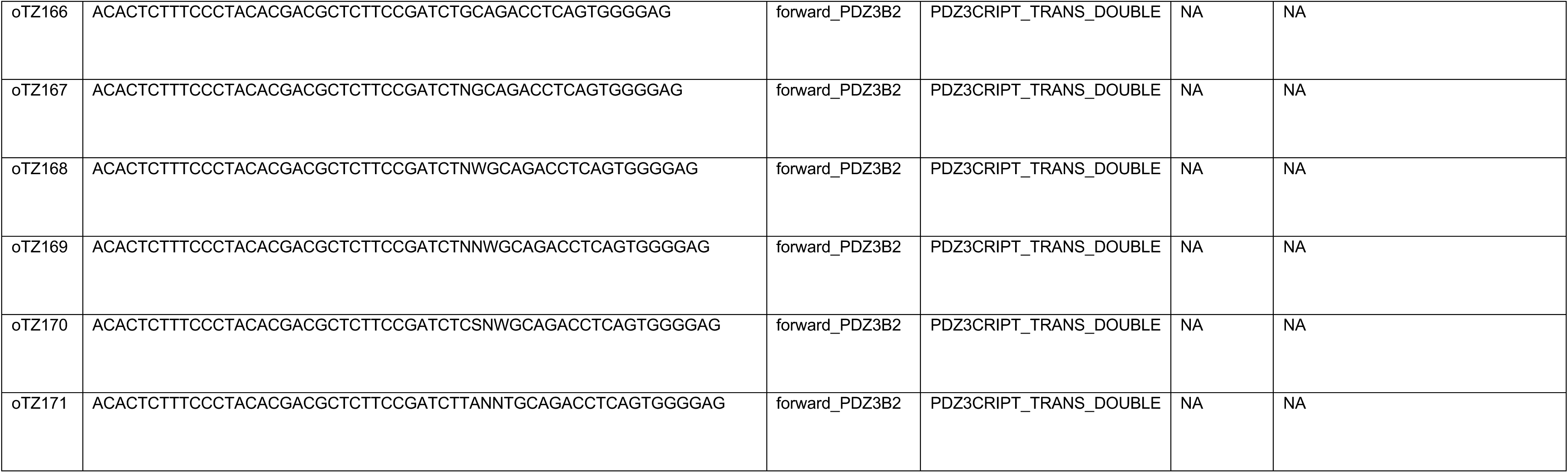
List of frameshifting oligonucleotides used for PCR1.

